# 4Pi MINFLUX arrangement maximizes spatio-temporal localization precision of fluorescence emitter

**DOI:** 10.1101/2023.10.30.564755

**Authors:** Julian D. Rickert, Marcus O. Held, Johann Engelhardt, Stefan W. Hell

## Abstract

We introduce MINFLUX localization with interferometric illumination through opposing objective lenses for maximizing the attainable precision in 3D-localization of single inelastic scatterers, such as fluorophores. Our 4Pi optical configuration employs three sequentially tilted counter-propagating beam pairs for illumination, each providing a narrow interference minimum of illumination intensity at the focal point. The localization precision is additionally improved by adding the inelastically scattered (fluorescence) photons collected through both objective lenses. Our 4Pi configuration yields the currently highest precision per detected photon among all localization schemes. Tracking gold nanoparticles as non-blinking inelastic scatterers rendered a position uncertainty < 0.4 nm^3^ in volume at a localization frequency of 2.9 kHz. We harnessed the record spatio-temporal precision of our 4Pi MINFLUX approach to examine the diffusion of single fluorophores and fluorescent nanobeads in solutions of sucrose in water, revealing local heterogeneities at the nanoscale. Our results show the applicability of 4Pi MINFLUX to study molecular nano-environments of diffusion and its potential for quantifying rapid movements of molecules in cells and other material composites.

## Introduction

The spatio-temporal precision of molecular tracking is generally limited by the information on the position of a molecule that can be retrieved in short time. In experiments using nanometer-sized fluorophores as molecular proxies, the maximum number of fluorescence photons that can be detected per unit time is the limiting factor for the achievable spatio-temporal precision. In principle, the fluorescence detection rate can be increased by applying a higher excitation intensity. At higher intensities, however, fluorophores transiently evade to non-emissive states or bleach. Increasing detector sensitivity or the transmissivity of the optical components is not an option either, since in a state-of-the-art optical setup these factors are already maximal. The only realistic way to improve molecular tracking is to design localization methods that augment the position information relayed by each detected photon. This is exactly the strategy that we pursue in this study.

In the most basic optical localization process, a measurement with precision *σ* is repeated photon by photon in order to refine the information about the position of the emitter. Statistically, the achievable precision is 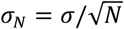, where *N* is the number of consecutive measurements (1). Most single fluorophore tracking experiments that use camera-based localization (2, 3) rely on such an identically repeated measurement process. Concretely, *N* photons are sequentially detected on a camera without any further modification of the measurement between individual detections. In each detection, the precision *σ* is governed by the diffraction of fluorescence light, which typically amounts to about 120-180 nm. Since the localization precision inevitably scales with 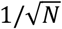, even under optimal optical conditions, more than 20,000 detected photons are needed to achieve *σ* < 1 nm (2). Even so, as fluorophores are dipole emitters whose emission is not isotropic right from the outset, the precision is slightly compromised by the temporal distribution of the fluorophore orientation in space (4).

MINFLUX localization (5-9) overcomes the 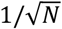 scaling by iteratively refining the position measurement gained from the preceding, inherently less precise localization. To avoid exclusive dependence on the diffraction of the fluorescence light, MINFLUX employs an illumination beam featuring a central point or line with vanishing intensity, i.e. a minimum. In iterative MINFLUX localization, this (nearly) zero intensity minimum is gradually moved closer to the emitter position. Due to the nearly quadratic curvature of the intensity around the minimum, the emission rate of the inelastic scatterer or fluorophore would scale accordingly. However, in the MINFLUX localization process the total power of the excitation beam is increased (with decreasing emitter-to-zero-distance) so that the emitter keeps experiencing the same (or similar) excitation intensity and the emission rate remains constant. At the same time, the decreasing emitter-to-zero distance enhances the positional information relayed by each detected photon. As a result, the localization precision scales by and large with *exp*(−*N*) (9).

This iterative localization eventually reaches a point where further refinement in terms of bringing the minimum closer to the fluorophore, and making the spatial interval *L* (5) smaller, is futile due to noise. In any case, the minimum *L* in which the emitter is to be expected is substantially smaller than the diffraction limit, with typical *L* values of 10-30 nm in the focal plane and 20-100 nm along the optic axis. At this point, the localization can still be refined by a standard localization scaling with 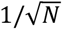. However, contrary to a standard localization, each detected photon now carries sub-diffraction position information due to the fact that the molecule is known to be located within a sub-diffraction interval *L*.

To allow the comparison of different localization methods, we introduce the single-photon efficiency 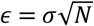 as a metric, where *σ* is the precision that is ultimately attained with *N* detected photons. Note that this metric does not depend on the emission rate and is equivalent to the precision achieved when detecting just one photon. In standard camera-based localization, *ϵ* is given by the precision at the diffraction limit, which is in the range of 120-180 nm; the value becomes larger in the presence of background. By contrast, in a MINFLUX measurement *ϵ* can be improved, i.e. made smaller in number, by more than an order of magnitude. In a MINFLUX system with non-zero (residual) excitation intensity or background *b* at the minimum, the achievable *ϵ* is proportional to 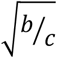 with *c* denoting the curvature of the profile around the intensity minimum (SI Section 2). Therefore, optimizing 3D MINFLUX localization calls for maximizing the curvature *c* while minimizing *b*, in all three dimensions. At first, these demands seem incompatible. While the curvature *c* can be increased by increasing the total power of the excitation beam, this measure concurrently increases the background *b*. At this point, a 4Pi optical arrangement (10) becomes highly relevant. By relying on two opposing lenses with counterpropagating interfering beams, a 4Pi setup readily provides an increase in curvature *c* of the interference intensity minimum along the optic axis by more than 200 % compared to any single-lens configuration (Fig. S4-6 and SI Section 3) and thus the most favorable condition for MINFLUX localization (11).

Guided by this reasoning we now demonstrate 3D localizations with nearly optimal beam curvatures attained by a changing pair of counterpropagating excitation beams in a 4Pi configuration. To identify the position in 3D, the beam pair is sequentially tilted along three different directions, effectively performing three independent 1D localizations. For symmetry reasons and simpler position evaluation, the three tilted axes have the same angle with respect to the symmetry (optic) axis of the two juxtaposed lenses. We show that this arrangement allows a precise localization of the 3D coordinate of an emitter. Furthermore, by collecting the photons with two high-NA (numerical aperture) objectives, the 4Pi configuration readily doubles the number of photon detections. Combined with the larger curvature, this photon doubling yields a more than 100-fold increase in MINFLUX tracking speed compared to popular camera-based localization techniques. In fact, the achieved spatio-temporal precision exceeds previously reported values, including those reported so far with MINFLUX. As a result, the system allows 3D measurements on fast diffusing single molecules and fluorescent beads with nanometer per millisecond spatio-temporal precision, revealing previously unobserved particle dynamics in viscous media.

### Setup

Our 4Pi configuration (Fig. 1A) entails a pupil inversion that rotates the position of the beam in the back focal plane of lens 1 by 180° with respect to the beam entering the sample through lens 2. Thus, the beams counterpropagate along the same axis. As the two beams are mutually coherent, the intensity minimum required for MINFLUX is rendered by destructive interference at the common focal point of the opposing lenses. The actual localization consists of three successive measurements, each performed along the separate axes x’,y’, and z’. Each axis is tilted by 41° with respect to the central (optical) axis of symmetry (Fig.1). The position in the (x,y,z)-laboratory space is obtained through an affine transformation from (x’,y’,z’)-space (SI Section 1.3). To allow for the 41° tilt, each beam fills the entrance pupil only partially.

**Figure 1.**
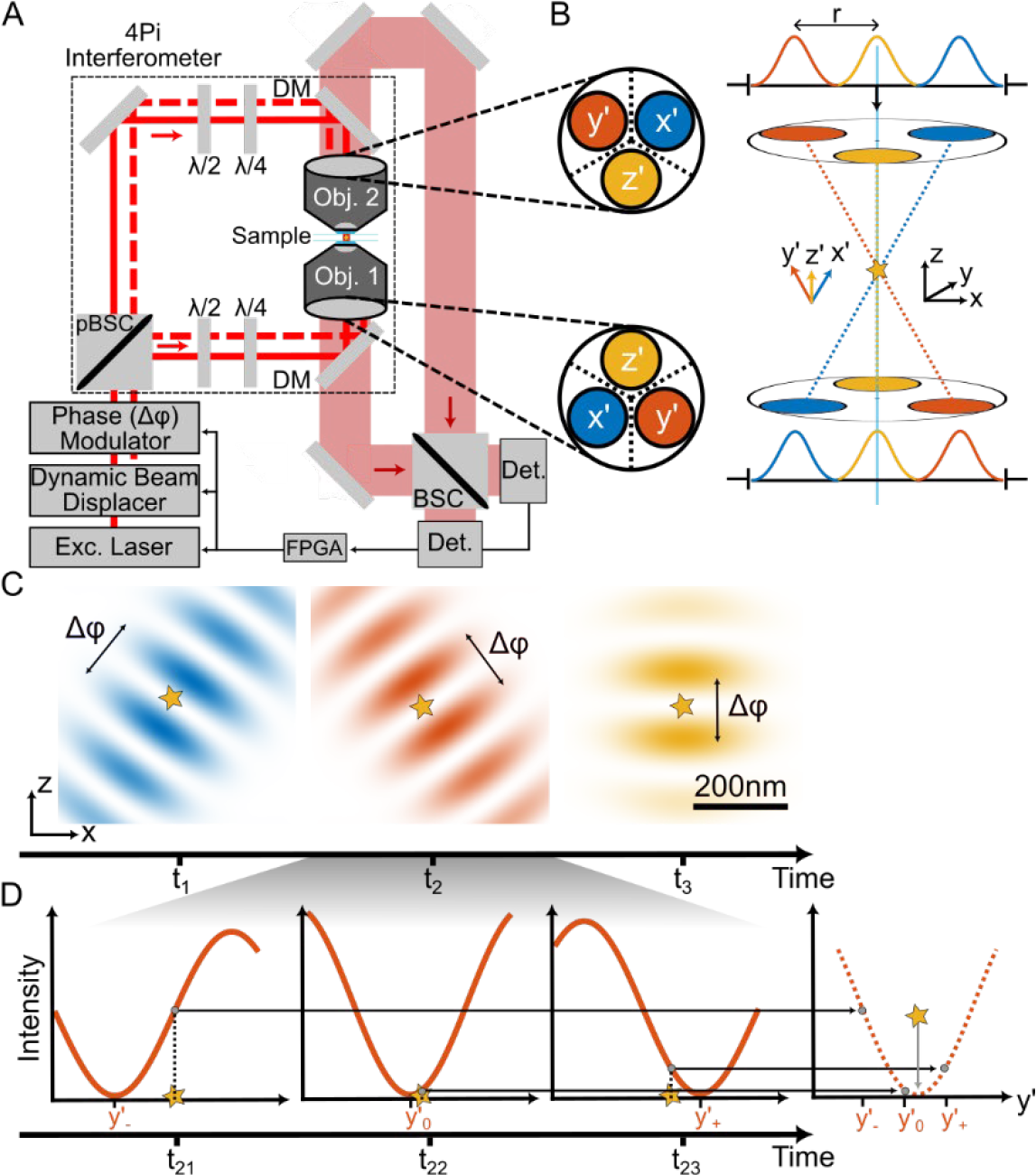
(**A**) Simplified layout of the 4Pi configuration with optically inverted entrance pupils of the two opposing objective lenses (Obj. 1, Obj. 2). The dynamic beam displacer shifts the position of the linearly polarized excitation beam from the laser in the entrance pupil of the lenses to three distinct positions (blue, red, yellow), yielding the measurement axes x’,y’, and z’ in the focal region. The dashed red line indicates two slightly offset partial beams, e.g. for y’, whereas the solid red line sketches the optic axis. The polarization of the laser beam is initially oriented at 45° to the interferometric plane and, after passing the beam displacer, partitioned by the polarizing beam splitter cube (pBSC) into orthogonally oriented (0° and 90°) beam pairs. The half-wave (λ/2) and quarter-wave (λ/4) plates in each interferometric arm rectify and adjust the polarization such that the counterpropagating partial beams interfere in the joint focal region of the lenses (either with linear or circular polarization). By shifting the relative phase Δφ between the two polarization directions, the phase modulator moves the intensity minimum along the applied tilted axis (x’,y’, or z’), enabling the iterative MINFLUX localization procedure along that axis. Note that the symmetry of the interferometer entails a point reflection of the beam through objective lens 1 with respect to that through lens 2, so that the two beams counterpropagate along the same axis. The fluorescence light collected by both lenses passes the dichroic mirrors (DM) and, after incoherent combination at the beam splitter cube (BSC), it is measured by two confocal detectors (Det). A field programmable gate array (FPGA) controls the laser, the beam displacer and the phase modulator; it also registers the signals from the two detectors. (**B**) Positions of the beams in the back focal plane for creating the x’,y’, and z’ axes for the MINFLUX measurement and relation of the pertinent coordinate system with the x,y, and z laboratory system; star indicates single emitter. (**C**) Projection of all three intensity distributions onto the xz-plane for sequential measurements. A phase difference Δφ introduced by the phase modulator places the local minimum to distinct positions along the relevant x’,y’, and z’ axes with the Δφ–induced track range denoted by the arrow. (**D**) MINFLUX localization procedure: fluorescence photons are collected for three positions of the excitation minimum within a range *L* around the unknown position of the emitter along one of the measurement axes (x’, y’, or z’). The resulting position on the respective axis is computed from the number of detected photons.

To tilt the counterpropagating beams, the juxtaposed position of the beams in the entrance pupils of the lenses is consecutively swapped by the same azimuthal angle of 120° (Fig. 1B). A dynamic beam displacer consisting of acousto-optical deflectors induces the position change (Fig. 1C). In fact, within the range tolerated by the entrance pupil, this approach allows any tilt of the standing wave and thus of the intensity minimum relative to the optical axis. Consequently, the three-axis measurement is readily performed dynamically. Since the aperture of the two beams is 3 -4 times smaller than that of the lens, the diameter of the intensity distribution of the standing wave is larger by the same factor than in a system with fully illuminated aperture (Fig. 1C). The curvature of the minima along the x’,y’, and z’ axis is improved by a factor of 1.45 compared to reported single lens illuminations), which is highly beneficial for locating the emitter by MINFLUX. Due to the 41° tilt of the (x’,y’,z’) coordinate system, the localization precision in the z direction is further increased, while the precision along the lateral dimensions is reduced. Overall this results in an increase in localization precision by a factor of 1.68 in z and 1.11 in x and y over a single-objective MINFLUX system using a line shaped local minimum (SI Section 3). The position of the intensity minimum along the axis of counterpropagation is controlled by a phase modulator acting as a phase shifter. The modulator electro-optically imprints a phase difference between the two polarization directions, which are separated at the polarizing beam splitter cube (pBSC) defining the entrance port of the 4Pi arrangement. For a single MINFLUX localization along one of the tilted measurement axes (x’,y’, or z’), the local illumination minimum probes the position of the emitter by counting the photons detected at three known positions of the minimum. These three positions are moved as closely as possible to the emitter along the individual axis (Fig. 1D). The position of the emitter along the axis is inferred from the number of detected photons.

## Results

Next we investigated the localization performance of our 4Pi MINFLUX setup using sequentially tilted standing waves. As 4Pi optical configurations are alignment sensitive, the beams were readjusted for each new sample. All measurements were taken along the same sequentially tilted axes positioned on a circle with radius r = 2 mm in the entrance pupil of the lenses. As in previous MINFLUX localizations, the three exposures per axis were equidistantly spaced and centered around the last emitter position estimate. Each tracking measurement started with a two-step zoom-in with a fixed distance L between the two outer positions of the minimum. Denoting the spatial interval where the fluorophore was expected to be located, the L-values for the zoom-in process were initially set to L_1_= 120 nm and subsequently to L_2_=80 nm.

### 3D-4Pi MINFLUX performance on single molecules

To determine the attainable single-photon efficiency *ϵ* of the system under standard experimental conditions, we prepared surface-immobilized ATTO647N fluorophores (SI Section 1.1.1) and continuously 3D-localized each of a total of 17 molecules until bleaching. After a two-step zoom-in process reducing L (see above), the distance between the two outer minimum positions was set to L_3_ = 40 nm with a duration of 300 μs per exposure. Following each 3D measurement using the tilted axes (x’,y’,z’), an additional measurement was taken along the optical axis (z) of the system in order to compare the signal-to-background ratio during localization with the outer beams with the signal-to-background ratio utilizing a central beam.

A representative trace of one of the fluorescent molecules observed (SI, Fig. S7) shows that the beam quality of the outer beam could be adjusted to a mean signal-to-background ratio of 4.1 compared to 6.2 for the central beam, indicating that the beam quality of the outer beams is controllable by the dynamic beam displacer. The position of a fluorophore was determined with a precision of 4.6 nm, 4.0 nm, and 1.6 nm in the x,y, and z direction, respectively, with only 62 detected photons per axis (SI, Fig. S7). The pertinent 3D single-photon efficiency is *ϵ* = 24 nm. The average single-photon efficiency for all 17 measured traces yields *ϵ* = 31 nm. For comparison, to our knowledge, the best single-photon efficiency in a classical single-molecule localization study was achieved on a 4Pi STORM microscope with *ϵ* = 160 nm (12), and the previous best single-photon efficiency in a 3D MINFLUX measurement was *ϵ* = 40 nm (13). Thus, the 4Pi microscope achieved the best 3D single-photon efficiency on single molecules to date.

Note that due to a skewed coordinate system, the precision along the optical axis is 51 % higher than the precision in the focal plane. A Cartesian coordinate system with almost 90° angles between each of the axes (and thus completely isotropic 3D precision) could possibly be realized only by placing the beams further out in the entrance pupil of the lens, albeit at the expense of the beam aperture and quality. We also observed that for 1D MINFLUX localizations along the optical (z) axis, the average single photon efficiency could be improved to *ϵ* = 6.3 nm (Fig. S9), owing to the optically favorable conditions offered by low aperture z-counterpropagating beams.

### 4Pi MINFLUX localization performance in 3D on idealized emitters

To test the 4Pi MINFLUX system without being affected by fluorophore blinking and bleaching, which both are not of fundamental relevance to the localization concept, we recorded the trajectory of inelastically scattering gold nanoparticles (SI Section 1.1.2). The constant stream of scattered photons was shifted to longer wavelengths compared to that of the laser and could thus be easily separated by appropriate optical filters. For this measurement, we set L_3_ = 20 nm during the tracking procedure and allotted 30 μs per exposure. Thus, we obtained a position measurement rate of 2.9 kHz for a point in 3D space. The trajectory of the nanoparticles was elicited by a controlled movement of the sample stage with an amplitude of 10 nm in x, y, and z. In this measurement (Fig. 2A), we achieved a signal-to-background ratio of 5.3, 2.7, 6.8 along the x, y, and z axes of the laboratory system (Fig. 2B), resulting in localization precisions (Fig. 2C) of 5.2 Å, 5.2 Å, and 3.3 Å, respectively. These values amount to a spatio-temporal precision of 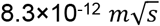 and single-photon efficiency *ϵ* = 9.4 nm in 3D localization, demonstrating the highest precision and efficiency among all 3D (MINFLUX) measurements to date.

**Figure 2.**
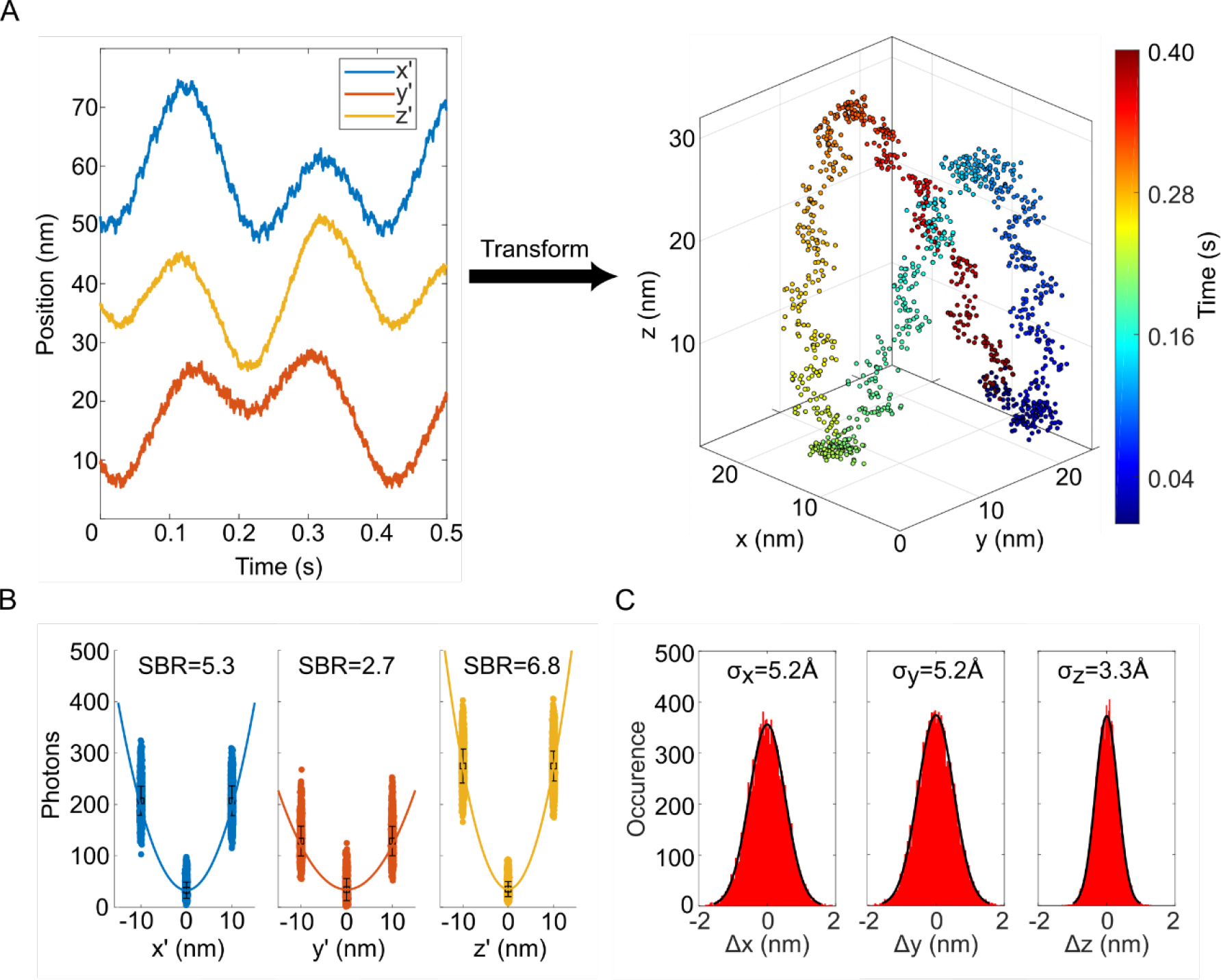
(**A**) Trajectory of an inelastically scattering gold nanoparticle (GNP) in the (x’,y’,z’) coordinate system in a 3D tracking measurement. The GNP was deliberately moved along a Lissajous figure trajectory using the sample stage. Coordinate transformation to the laboratory system (x,y,z) reveals the actual 3D trajectory. (**B**) Distribution of photon numbers among the three exposures used for MINFLUX localization, shown for localization along the x’,y’ and z’ axis and with the pertinent signal-to-background ratio. (**C**) Histograms of position differences between two consecutive measurements in the laboratory system demonstrating short-term localization precision at Ångström scale at a 3D detection rate of 2.9 *kHz*.

### Nanometer-scale 3D MINFLUX diffusion tracking

In a simple application we took advantage of the performance of 3D-4Pi MINFLUX to gain insight into 3D diffusion at the nanoscale, specifically of single-particle tracking of small fluorescent beads and single fluorophores in solution (SI Section 1.1.3). In general, diffusion is described by the diffusion coefficient *D*, which is related to the movement of a particle as *D*(*t*) = MSD(*t*)*/*(2*n*_*d*_*t*) (14), where MSD(*t*) is the mean-squared displacement after time *t* and *n*_*d*_ is the dimensionality of the space in which the particle diffuses. The diffusion coefficient can take different forms depending on the type of diffusion, e.g. hop diffusion for obstructed movements or free diffusion that is purely stochastic. In the case of thermally driven stochastic motion in a liquid, the diffusion coefficient is given by the Stokes-Einstein relation *D* = *k*_*B*_ *T/*(6*πηR*_0_). Here, *k*_*B*_ is the Boltzmann constant, whereas *T, η* and *R*_0_ are the absolute temperature, the macroscopic viscosity of the fluid and the hydrodynamic radius of the diffusing particle, respectively. As particle radii become smaller the diffusion coefficient increases, leading to faster movement. Therefore, the study of the diffusion of small particles inherently requires high spatio-temporal precision.

We used oversaturated sucrose-water solutions as a model system for the diffusing fluorophores and beads, because these solutions have viscosities of several pascal-seconds, which is in the range of viscosities reported for cellular environments (15, 16). We chose LD655 as the fluorophore because it exhibits good photostability in viscous media due to its cyclooctatetraene group preventing the formation of reactive oxygen species (17). This aspect is particularly beneficial in environments where diffusive buffer systems are not applicable.

Our 3D MINFLUX tracking measurements relied on L_3_ = 80 nm at 100 μs per each of the 9 exposures, resulting in a position acquisition rate of about 1 kHz. For each trace, we calculated the mean squared displacement of the movement. Tracks of single LD655 molecules and of 27 nm diameter fluorescent beads (Fig. 3) were recorded. Surprisingly, a subset of particles showed hop-like instead of stochastic diffusion, suggesting a previously unreported environment in an oversaturated sucrose-water solution (SI Section 1.5). Hence, we performed a maximum likelihood classification in which each individual trace was examined to determine whether it exhibited stochastic or hop-like diffusion properties. The validity of the analysis routine was initially tested on Monte-Carlo simulated data (SI Section 1.6). For the LD655 molecules, we evaluated 72 traces with a duration of *t* = 49 ms each. The average spatial displacements between consecutive positions in the laboratory system were 10.6 nm, 11.8 nm, 7.0 nm in x, y, and z-axis respectively, resulting in a 3D displacement of 9.6 nm in length. Similarly, we analyzed 198 fluorescent beads traces of *t* = 195 ms duration, where we found corresponding average spatial displacements of 9.2 nm, 11.7-nm, 6.0 nm, resulting in a 3D displacement of 8.6 nm.

**Figure 3.**
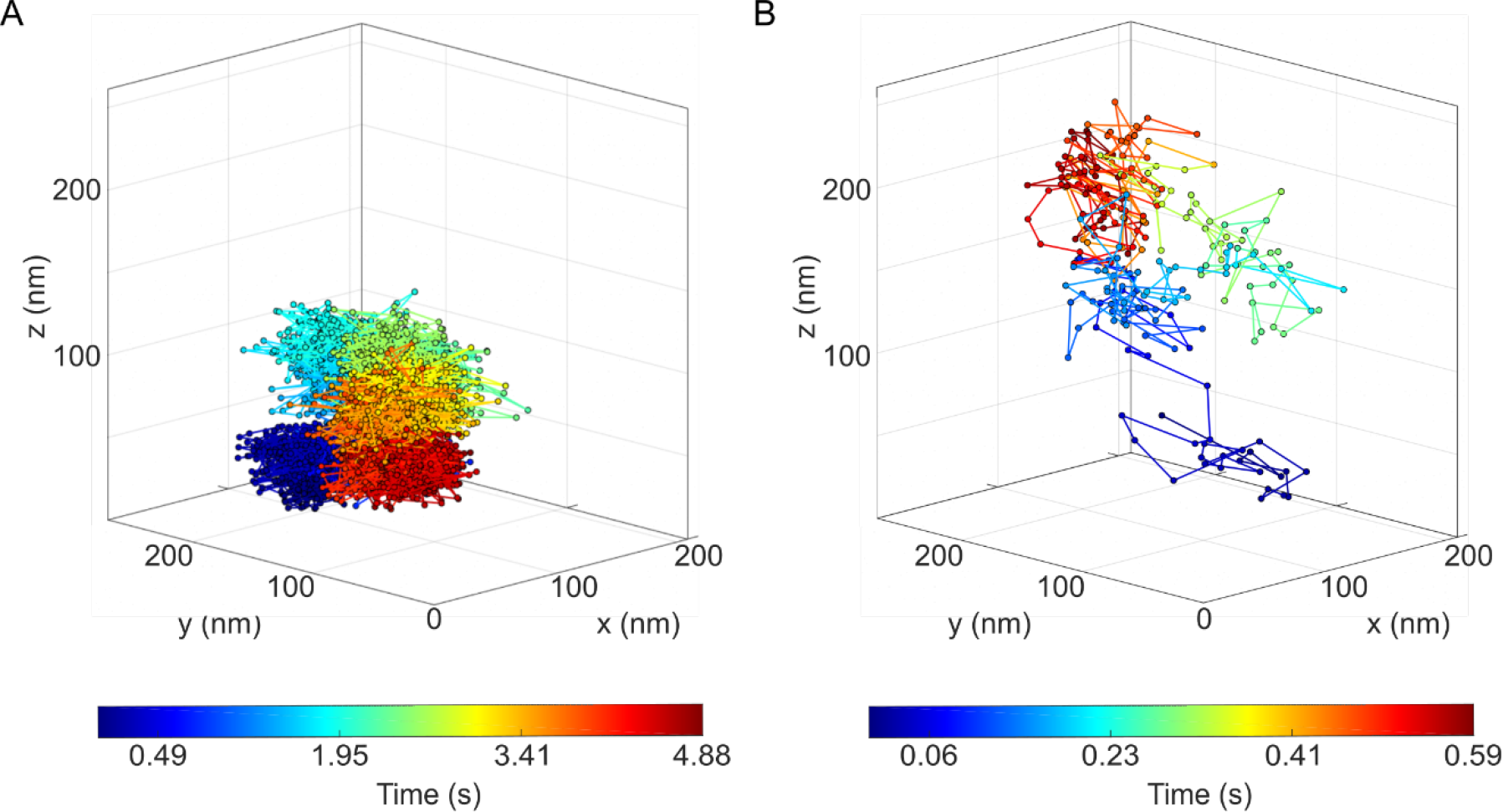
**(A)** Example 3D trajectory of a 27 nm diameter fluorescent bead diffusing in a sucrose solution, obtained by a 4Pi MINFLUX setup. A total of 4383 positions were obtained over a period of 4.88 seconds; the time for a single measurement was 975 μ*s*. **(B)** Example 3D trajectory of a single LD655 fluorescent molecule diffusing in a sucrose solution obtained in a 4Pi MINFLUX setup. A total of 241 positions were obtained over a period of 0.59 seconds; the time for a single position measurement was 975 μ*s*.

Equating the consecutive displacements with the localization precision (as a conservative estimate of its upper bound) yields an average single-photon efficiency of *ϵ* = 65 nm for the beads and *ϵ* =79 nm for the single fluorophores. Since *ϵ* is still below the diffraction limit, we can conclude that any classical localization method that does not refine the measurement during localization, as is the case in MINFLUX, would be limited in recording the particle movement. Indeed, deviations between observed and expected diffusion have been reported for popular camera-based particle tracking (18). While several computational approaches have been employed to mitigate this effect (19, 20), these methods are limited to testing specific diffusion models. General improvements can only be achieved by a measurement scheme that increases the information provided by the detected photons.

The classification shows that for the single LD655 fluorophores (Fig. 4B), 29% of the traces are classified as hop-like diffusion, which implies a separation of diffusion into two time scales, whereas for the fluorescent beads, 89% of the diffusion traces are hop-like (Fig. 4A). According to the analysis of the simulations (SI Section 1.4), this finding suggests that the single fluorophores exhibit different diffusion modes, while the beads mostly exhibit hop-like diffusion.

**Figure 4.**
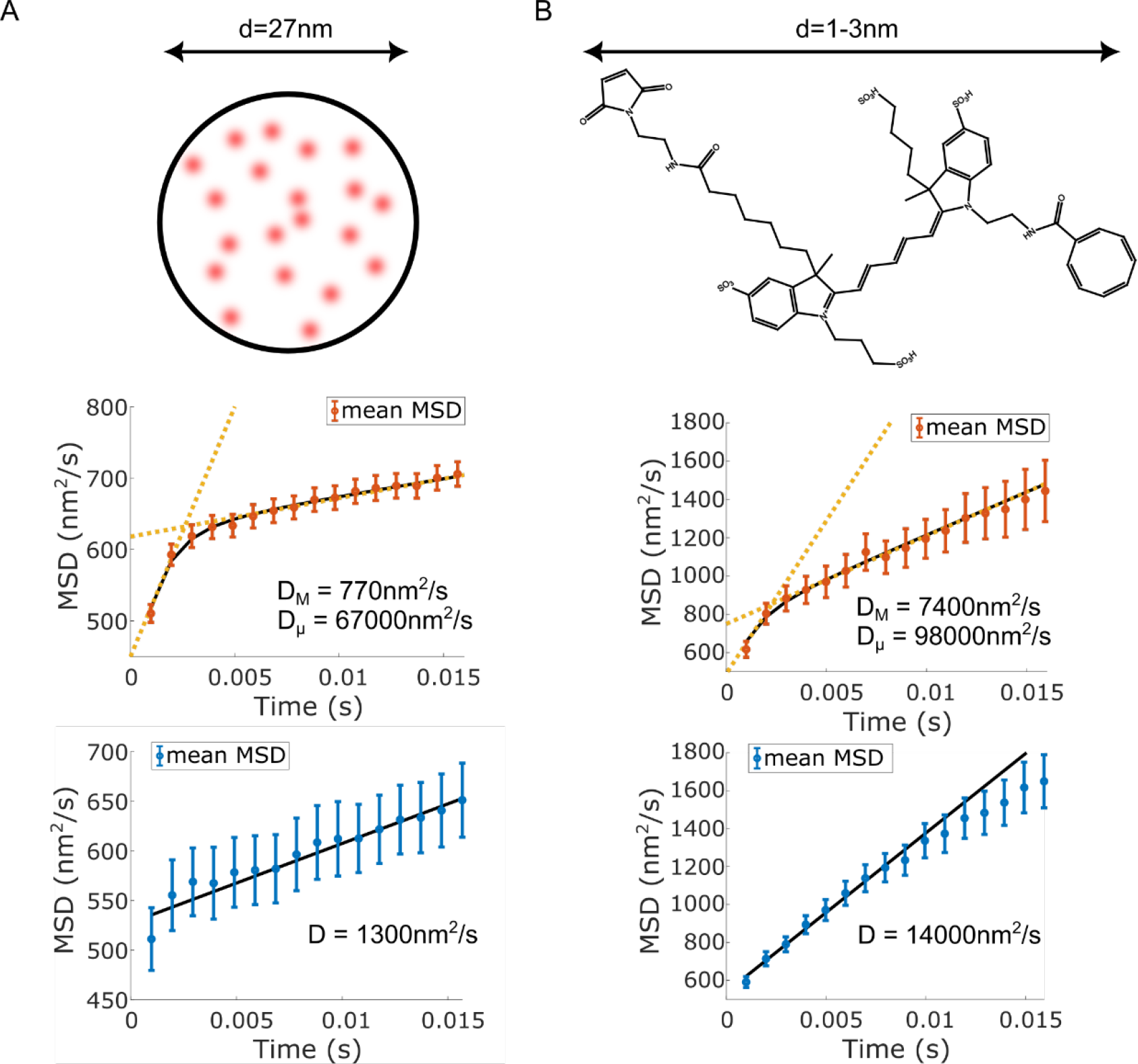
(**A**) Sketch of the fluorophore-stained bead used and result of the evaluation of 198 bead traces, which were classified into hop-like (red, ∼89%) and stochastic (blue, ∼11%) diffusion. Fits to the mean of the bead mean squared displacements (MSD) classified as hop/stochastic diffusion (middle/bottom). The difference between short-term and long-term diffusion is apparent from the difference in diffusion coefficients *D*_*M*_ and *D*_*μ*_ by about 87-fold, as indicated by the yellow dashed trend lines. The deviation of the first point in the stochastic diffusion data is because some traces showing hop-like diffusion are misclassified as stochastic diffusion by our analysis routine (see SI Section 1.6). (**B**) Structure and approximate size of the diffusing LD655 fluorophore and result of the evaluation of 72 single fluorophore traces, which were classified into hop-like (red) and stochastic (blue) diffusion yielding a partition into 29 % and 71 %, respectively. Fits to the mean of all fluorophores’ mean squared displacements (MSD) classified as hop/stochastic diffusion (middle/bottom). The different time scales are highlighted by the different trend lines (yellow dashed) for short-term and long-term diffusion. Note the ∼13-fold disparity in diffusion coefficients *D*_*M*_ and *D*_μ_ Diffusion on the nanoscale is much faster than diffusion on longer length scales.

The measurements also show that the diffusion of smaller particles is slower on macroscopic scales (Fig. 4) and scales inversely with particle diameter, as predicted by the Stokes-Einstein relation. In addition, the classification analysis reveals hop-like diffusion for some of the observed particles with a separation of time scales around 2-3 ms. Some particles undergo hop-like diffusion while others show stochastic diffusion, highlighting that diffusion at the nanoscale depends on the nano-environment. The inhomogeneity of the nano-environment is also evident in the variation of the fitted parameters (Fig. S8). A possible explanation for this behavior is that the sucrose molecules may undergo oligomerization as has also been reported for other carbohydrates (21, 22), making it more difficult for larger particles to penetrate the solution.

## Conclusion and Outlook

We have introduced a 4Pi setup with mutually inverted and partially filled entrance pupils for 3D MINFLUX localization. The setup allows the creation of a standing wave that is quickly tilted in three different orientations with respect to the optical axis. As the interfering beams are of reduced aperture and counterpropagating head-on, the interference intensity minima attain the virtually highest possible curvature around the zero intensity point. Thus, our 4Pi MINFLUX approach provides highly favorable conditions for performing 3D localization with maximal spatio-temporal precision. Using the definition of the single-photon efficiency 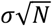 and the average emission rate *R* = *N*/*t*, the spatio-temporal precision (STP) of the emitter localization can be written as 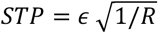. For single fluorophores with uninterrupted emission, we attained an *STS* of 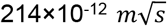, whereas for constantly inelastically scattering gold beads we achieved an 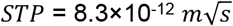.

As our 4Pi MINFLUX approach effectively maximizes *ϵ* and the efficiency of photon detection, future improvements should primarily concentrate on improving the inelastic scattering properties of the observed emitters, particularly the fluorescence emission rate. Remaining limiting factors are aberrations of the optical system and the inevitable background signal. Since aberrations cause a (non-zero) depth of the excitation intensity minimum, these factors are intertwined. Specifically, for the MINFLUX localization process the fluorescence generated at the minimum adds up to the background arising from stray light or detector dark counts to form the total background *b*. Aberrations also reduce the curvature of the illumination profile *c* and thus the achievable localization precision. In a 4Pi configuration, careful calibration and beam path optimization are required to avoid non-negligible asymmetries between the interfering beams. As each sample is slightly different, we continuously refined the beam paths using the dynamic beam displacer. In addition, since the sample itself is part of the optical path, its refractive index should be as close as possible to that of the immersion medium.

In conclusion, the implementation of a 4Pi arrangement enables the most precise 3D tracking of fluorophore positions so far. Specifically, diffusion measurements on single fluorophores and fluorescent beads showed that 4Pi MINFLUX records the 3D movement of inelastically scattering emitters with hitherto unattained spatio-temporal precision. The use of non-blinking, bright emitters allowed us to demonstrate a record 3D spatio-temporal precision of *σ*_3*D*_= 4.5 Å per 345 μs. Therefore, 4Pi MINFLUX has the conceptual and practical power to improve the observation of nanoparticle dynamics in polymer nanocomposites (23-25), the molecular transport processes in cells (26), and of other 3D dislocations or movements of particles where highest spatio-temporal resolution is of paramount importance. Finally we note that, although our paper concentrated on tracking, in conjunction with sequential on-off switching of single fluorophores our 4Pi MINFLUX system should readily provide 3D images with record localization precision (per detected photon) and resolution at the molecular scale.

## Author Contributions

J.D.R., M.O.H., J.E., and S.W.H. designed research; J.D.R., M.O.H., and J.E. performed research; J.D.R. analyzed data; S.W.H. supervised the project; and J.D.R., J.E., and S.W.H. wrote the paper.

## Materials and Methods

A detailed description of the optical setup, the sample preparation, and the data analysis pipeline can be found in the Supplements.

## Acknowledgments

We thank Michael Dasbach for his help with the preparation of the diffusion samples. We thank Jade Cottam Jones and Steffen J. Sahl for helpful discussions and critical reading of the manuscript. This research was conducted within the Max Planck School Matter to Life supported by the German Federal Ministry of Education and Research (BMBF) in collaboration with the Max Planck Society.

