## Supplements for "4Pi MINFLUX arrangement maximizes spatio-temporal localization precision of fluorescence emitter"

*Supporting Information for*
### 4Pi MINFLUX arrangement maximizes spatio-temporal localization precision of fluorescence emitter

Julian D. Rickert[a,b], Marcus O. Held[a,1], Johann Engelhardt[a], Stefan W. Hell[a,c,*]

[a]Department of Optical Nanoscopy, Max Planck Institute for Medical Research, 69120 Heidelberg, Germany; [b]Max Planck School Matter to Life; [c]Department of NanoBiophotonics, Max Planck Institute for Multidisciplinary Sciences, 37077 Göttingen, Germany

[1]Present address: Molecular Microscopy and Spectroscopy, Istituto Italiano di Tecnologia, 16152 Genoa, Italy

[*]To whom correspondence should be addressed.

**Contents**

1. Materials and Methods
   - 1.1. Sample Preparation
     - 1.1.1. Preparation of ATTO647N samples
     - 1.1.2. Preparation of gold nanoparticle samples
     - 1.1.3. Preparation of samples for diffusion measurements
   - 1.2. Detailed description of 3D-4Pi MINFLUX microscope
     - 1.2.1. Laser module
     - 1.2.2. Beam displacement module
     - 1.2.3. Phase scanner module
     - 1.2.4. 4Pi interferometer
     - 1.2.5. Diagnostics module
     - 1.2.6. Detection module
   - 1.3. Transformation from the (x’,y’,z’) system into the laboratory system
   - 1.4. Measurement process
   - 1.5. Diffusion: Theoretical background
   - 1.6. Diffusion simulation and analysis routine
     - 1.6.1. Simulation
     - 1.6.2. Data analysis pipeline
     - 1.6.3. Data fitting
     - 1.6.4. Test of analysis with simulated data
2. Precision limit in MINFLUX nanoscopy
3. Precision limit in 3D-4Pi MINFLUX
   - 3.1. Comparison between single objective and dual objective MINFLUX

### 1. Materials and Methods

#### 1.1. Sample preparation

All 4Pi samples consist of one 18 mm, and one 15 mm diameter coverslip (170 µm, No 1.5H, Paul Marienfeld GmbH & Co. KG, Lauda-Königshofen, Germany) sandwiched together and sealed with either nail polish (RIVAL loves me Care Power Hardener, Dirk Rossmann GmbH) or two-component glue (2-K-Epoxidkleber Sofortfest, UHU, Bühl, Germany).
Before a coverslip is used for sample preparation, its thickness is checked to be within $\pm$ 3 µm of the average thickness of 170 µm. This step is taken to ensure equal beam paths for the bottom and top illumination beams. The coverslip sandwich is mounted on an aluminum ring manufactured in our mechanical workshop to fit precisely on the sample stage. All coverslips are washed in acetone and ultrapure water and then plasma cleaned (TePla 300 Semi-Auto Plasma Processor) for 5 minutes under 0.5 mbar of $O_2$ at a power of 200 W to remove contaminants from the surface.
The sample is mounted on the top of the lower 18 mm diameter coverslip.
This coverslip has an additional aluminum quarter mirror printed onto its upper surface to create a reflection from the illumination laser, which is used to check the two objectives for spherical aberrations and to minimize aberrations with the objectives' correction collars. In addition, residual aberrations occur mainly in the upper illumination path due to mounting media that are not perfectly index-matched. To minimize this effect, the sample thickness is adjusted to less than 20 µm if the index cannot be exactly matched.
For each sample, gold nanoparticles (GNP-bt, 40 nm Biotin Gold Nanoparticles, Cyto-diagnostics Inc, Burlington, Canada) are immobilized on the surface to facilitate the alignment procedure of the 4Pi MINFLUX system.
To functionalize the surface of the coverslip, the surface is coated with PLL-PEG-bt (PLL: Poly-L-Lysin, PEG: Polyethylene Glycol, bt: biotin) and then NVD (Neutravidin) is added. PLL-PEG binds to the glass surface to prevent the sample from sticking. The biotin is used to functionalize the coverslip so that the sample can adhere to it. NVD binds to some of the biotin binding sites to dilute the available sites for the sample to be added. The entire procedure consists of applying 50 µL of PLL-PEG-bt (0.2 mg/mL) with 1%Vol Tween(P20) for 15 min followed by 50 µl of NVD (1 mg/mL diluted 1:100 in phosphate buffered saline (PBS)) for 5 minutes. Between each step, the coverslip is washed with 1 ml of PBS and dried under nitrogen flow.

##### *1.1.1. Preparation of ATTO647N samples*

The ATTO647N single-molecule samples were prepared by plasma cleaning and functionalization of coverslips as described in SI Section 1.1. The coverslip was incubated with 50 µL of a mixture of Atto647N-bt and GNP-bt for 5 minutes. GNP-bt was diluted 1:100 from stock in PBS, and ATTO647N-bt was added to the mixture at a dilution of 100 pM. After 5 minutes of incubation, the coverslip was washed with 1 ml of PBS. Mounting medium consisted of PBS with 1% VectaCell (CB-1000-2, Vector Laboratories Inc., Burlingame, USA) added to it. VectaCell contains Trolox, which reduces the formation of reactive oxygen species.

##### *1.1.2. Preparation of gold nanoparticle samples*

Plasma cleaned and functionalized coverslips (see SI Section 1.1.) were used to prepare GNP samples. The coverslip was incubated with 50µl of a mixture of PBS with GNP-bt diluted 1:100 from stock for 5 minutes. After 5 minutes of incubation, the coverslip was washed with 1 ml of PBS. The mounting medium was 2,2'-thiodiethanol (TDE).

##### *1.1.3. Preparation of samples for diffusion measurements*

Oversaturated sugar solutions containing different fluorescent markers were prepared on a chemical bench before being mounted on a coverslip.
The solutions had a sucrose (S7903, Sigma-Aldrich, Merck, Darmstadt, Germany) to ultrapure $H_2O$ ratio of w/w 83/17, resulting in a viscosity on the order of η = 10-100 Pas and a refractive index of approximately 1.52, corresponding to the refractive index of the oil immersion. Fluorescent markers were added to the mixture at the desired concentration. The exact mass fractions of the two components were 7.5 g of $H_2O$ and 36.6 g of sucrose.
For single fluorophore experiments, we added 130 aM LD655 (LD655-NHS, Lumidyne Technologies, Brooklyn, US) diluted in 1 µl ultrapure $H_2O$ to the sugar solution. LD655 was selected for its high fluorescence rate in viscous media compared to other single molecules because it does not rely on an external buffer system to avoid long-lived dark states. This is achieved by a cyclooctatetraene (COT) coupled to the fluorophore, which prevents the formation of reactive oxygen species (Ref. 1).

For the experiments with fluorescent nanobeads, Crimson beads (FluoSpheres Carboxylate modified microspheres (Crimson), Sigma-Aldrich, Merck, Darmstadt, Germany) with a diameter of 27 nm were used. A solution was obtained by mixing the ultrapure $H_2O$ with 1 µl of the Crimson beads stock solution.

The ingredients were mixed in a 50 ml flask mounted to a reflux apparatus to minimize water loss and ensure reproducibility. The flask was heated to 120°C while stirring with a stirring rod at 550 rotations per minute. After 1 hour, the heating was turned off, and the resulting viscous liquid was immediately poured into a conical plastic tube placed in an ice bath. This was done to prevent spontaneous crystallization, which is more likely at higher temperatures. The solutions were stored in the refrigerator at 4°C for several days.

Before mounting the sugar solution, the coverslip was functionalized according to SI Section 1.1. Then 50 µl GNP 1:100 from stock in PBS were incubated on the surface for 5 minutes. The GNP were washed with 1 ml PBS and the coverslip was dried under nitrogen flow. A small amount of the sugar solution was then added to a 15 mm coverslip, which was then placed on top of the GNP-functionalized coverslip. After that, the sandwich was wrapped in a layer of tissue and pressed in a vise. The last step ensured a homogeneous distribution of the sugar solution and kept the sample thickness as low as possible.

#### 1.2. Detailed description of 3D-4Pi MINFLUX microscope

The whole microscope is placed on an air-cushioned optical table (I-2000 isolators and RS4000 plate, Newport Spectra-Physics GmbH, Darmstadt, Germany) to decouple it from the vibrations of the building. Furthermore, we build an aluminum frame (30 mm profile system, MayTec Aluminium Systemtechnik GmbH, Olching, Germany) around the microscope, which contains two layers of acoustic foam (Pur Skin, SONATECH GmbH + Co. KG, Ungerhausen, Germany). This is done to shield the microscope from airflow and acoustic noise. All lasers are mounted on another optical table to avoid vibrations from cooling fans and heat transfer to the MINFLUX microscope. In addition, most of the electronics and the PC are placed on an additional frame that is coupled to the wall directly above the optical table. This allows the use of short cables and also decouples the microscope from vibrations. In addition, parts of the microscope are separated by darkening walls to reduce airflow inside the microscope and ensure temperature stability.

The optical system is shown in Fig. S1. The whole system can be divided into several modules: A laser module, the beam displacement module, the phase scanner module, the 4Pi interferometer, a diagnostics module, and a detection module.

##### *1.2.1. Laser module*

The laser module is mounted on a standalone optical table. The excitation laser light is generated by a single-frequency laser with a wavelength of 640 nm (Bolero, Cobolt AB, Solda, Sweden). The beam passes through an acousto-optical tunable filter (AOTF,PCAOM-VI, Crystal Technology Inc., Palo Alto, USA), which modulates the intensity. The excitation light is then coupled into a polarization-maintaining single mode fiber (SMF, P3-630PM-FC-10, Thorlabs Inc., Newton, USA) via a fiber collimator (L1, 60FC-4-RGBV11-47, Schäfter + Kirchhoff GmbH, Hamburg, Germany). A half-wave plate (λ/2, half-wave plate, B Halle, Berlin, Germany) rectifies the polarization of the beam to the transmission direction of the fiber.

##### *1.2.2. Beam displacement module*

Before entering the beam displacement module, the excitation light is collimated by a fiber collimator (L1) before passing through a half-wave plate (λ/2, half-wave plate, B Halle, Berlin, Germany) to set the polarization perpendicular to the optical table. A laser line filter (F1, ZET405/488/561/640xv2, AHF Analysentechnik AG, Tübingen, Germany) then spectrally filters the beam to remove light frequencies which that may be generated in the optical fiber due to phase fluctuations. The beam is then passed through a telescope consisting of two lenses (L2, f = 60 mm and L3, f = 30 mm). The telescope ensures that the beam pattern generated in the beam displacement module can be accommodated by the crystal optics in the phase scanner module. A Glen Laser polarizer (GL, GL10-A, Thorlabs Inc., Newton, US) cleans the polarization before the beam enters the pair of 2D acousto-optical deflectors (2D-AODs, AA.DTSXY-400-532(450.650)-011, Pegasus Optik GmbH, Wallenhorst, Germany). The deflectors displace the beam along the two directions perpendicular to the axis of propagation.

##### 1.2.3. Phase scanner module

A GL cleans the polarization and sets it perpendicular to the optical table. Then, a motorized half-wave plate (motλ/2, half-wave plate, B Halle, Berlin, Germany) mounted on a motorized precision rotation stage (PRM1/MZ8, Thorlabs Inc., Newton, US) is used to tilt the polarization by about 45°. This ensures equal intensities in the two interferometer beam paths inside the 4Pi interferometer. The beam then passes through two rubidium titanyl phosphate (RTP) crystals (EOM1 and EOM2, 8 × 8 × 25 mm custom build, CRISTAL LASER, Messein, France) which act as phase modulators.

##### 1.2.4. 4Pi interferometer

After the phase scanner module, the beam passes through a telescope (L3 and L2). This serves two purposes. First, it increases the displacement of the beam displacement module by a factor of two. Second, it increases the beam diameter by a factor of two, ensuring collimated beams inside the 4Pi interferometer. Behind the telescope, the beam is split into two paths by a polarizing beam splitter cube (PBSC, PBS201, Thorlabs Inc., Newton, US). After the PBSC, both beams pass through identical optical elements. This ensures symmetry between the two beams, which generate an interference pattern between the objectives at the position of the sample. Additional mirror reflections in one of the two paths create a relative beam inversion between the two paths. In addition, one of the mirrors in the lower path can be moved by a tip/tilt piezo scanner (S-316, Physik Instrumente (PI) GmbH & Co. KG, Karlsruhe, Germany) to obtain confocal images through the lower objective. Inside the interferometer, both beams are guided through a Glen Laser polarizer (GL) and a pair of motorized half-wave and quarter-wave plates (motλ/2 and motλ/4). This provides complete control over the polarization state of the two beams in the pupil plane of their respective objectives. Each interferometer arm contains a mechanical beam block (mechBB) that is used to switch between illumination through one objective and illumination through both objectives. The illumination light and the fluorescence light are split at a dichroic beam splitter (DM, ET435/510/595/705, AHF Analysentechnik AG, Tübingen, Germany). Depending on the sample, a pair of oil immersion objectives (HC PL APO 100x/1.44 OIL CORR CS, Leica Microsystems GmbH, Wetzlar, Germany) and a pair of water immersion objectives (Leica HC PL APO STEDwhite 86x/1.20 W motCORR, Leica Microsystems GmbH, Wetzlar, Germany) are used. Water immersion objectives were used for the ATTO647N measurements, while oil immersion objectives were used for all other measurements. The lower objective can be moved in the lateral plane by two piezo motors, while the upper objective can be moved in the axial direction by one piezo motor (Piezo LEGS® Linear Twin - C 450N, PiezoMotor Uppsala AB, Uppsala, Sweden). The sample itself is mounted on a customized sample holder, which is attached to the sample stage (Smarpod 110.45.2-d-sc-149, SmarAct GmbH, Oldenburg, Germany) via magnets. The position of the sample can be tracked in the laboratory system using an external interferometer (Picoscale, SmarAct GmbH, Oldenburg, Germany).

##### 1.2.5. Diagnostics module

Approximately one percent of the illumination light passes through the two DMs and into the diagnostics module. The two beams are combined on a beam-splitter cube (BSC, BS010, Thorlabs Inc., Newton, US). At a second BSC, the beams are split into two paths. The first beam passes through a lens (L4, 50 mm) and is imaged on a camera (Cam 1, daA2500-14um, Basler AG, Ahrensburg, Germany) in the focal spot of L4. The second beam is directed to a second camera (Cam 2, acA1920-155um, Basler AG, Ahrensburg, Germany) mounted in a plane conjugate to the pupil planes of the two objectives. The spots of the different beams used for a three-dimensional MINFLUX measurement are monitored on these cameras to allow referencing.

##### 1.2.6. Detection module

The fluorescence light from the two objectives is combined on a BSC. The two output ports of the BSC are each followed by a confocal detection path. Each of the detection paths contains a motorized pinhole (motPH, MPH16-A, Thorlabs Inc., Newton, US) to which a lens (L7, 75 mm) is attached. One of the detection paths has a lens (L5, 160 mm) with a focal length of 160 mm, forming a telescope with L7. This detection path consists of a single avalanche photodiode (APD1, SPCM-AQRH-13-FC, Excelitas Technologies, Waltham, MA, USA) with two filters (F2 and F3) in front of it. The filters block residual light from the excitation source. They can be exchanged depending on the wavelength used for the measurement. A lens (L4, 40 mm) focuses the fluorescence light onto the APD. The second detection beam path has a lens (L6, 200 mm) mounted in front of the motorized pinhole. Behind the pinhole, the detection light is spectrally split into two separate detection paths. The wavelengths around 700 nm are transmitted through the two bandpass filters F4 (FF01-698/70-25, Semrock Inc., Rochester, US). The transmitted light is focused by L4 onto APD2 (same model as APD1). The light deflected by F4 is directed to a photomultiplier tube (PMT, H9305-03, Hamamatsu Photonics K.K., Hamamatsu City, Japan). This PMT collects the residual laser light transmitted by the dichroic mirrors. It is used for alignment.

##### 1.3. Transformation from the (x',y',z') system into the laboratory system

The positions measured along the set of beam axes of the illumination beams are transformed into the laboratory system by an affine transformation that is determined for each sample. The transformation matrix is obtained by performing a MINFLUX measurement of a moving emitter along the desired beam axes while simultaneously determining the position of the sample in the laboratory system.

We measure the position in the laboratory system with an external interferometer mounted on a custom-made mount shown in Fig. S2. This mount is attached to a fixed part of the stage and houses three fibers that originate from a PicoScale interferometer (SmarAct GmbH, Oldenburg, Germany). Each of the fibers points to a single side of a mirrored cube attached to the sample holder. The three fibers are aligned so that an infrared laser is reflected back into the fiber, effectively forming a Michelson interferometer with a reference mirror mounted inside the fiber head.

To find the affine transformation $T_{\mathrm{MINFLUX,Lab}}$ from the MINFLUX beam system to the laboratory system, the microscope is aligned, and a gold nanoparticle (GNP) is brought into focus. Then, we induce a motion of the GNP by moving the stage successively in x, y, and z. The movements are square oscillations with a frequency of 3 Hz and an amplitude of 100 nm in the lateral direction and 50 nm in the axial direction. While the stage is moving, we perform a MINFLUX measurement along the desired coordinate axes and a measurement of the stage position using the PicoScale interferometer. The amplitudes of both measurements are extracted with a custom written MATLAB code. From these amplitudes, the transformations from the stage system to the MINFLUX system $T_{\mathrm{Stage,MINFLUX}}$ and from the stage system to the laboratory system $T_{\mathrm{Stage,Lab}}$ are obtained. Finally, from these two transformations, we calculate the direct transformation from the MINFLUX system to the laboratory system $T_{\mathrm{MINFLUX,Lab}} = \left(T_{\mathrm{Stage,Lab}}^{-1} T_{\mathrm{Stage,MINFLUX}}\right)^{-1}$.

Note that the set of axes for the MINFLUX measurement is arbitrary, as long as at least three individual axes are addressed. The calibration measurement described above is always performed for three beams, which are equidistantly placed on a circle, and an additional beam passing through the center of the objective.

##### 1.4. Measurement Process

The beams used for all of the measurements were positioned on a circle with radius r = 2 mm in the back focal plane of the objectives and had a FWHM = 1.5 mm. Each measurement process started with a two-step zoom-in with a fixed distance L between the two outer minimum positions. The L-values for the zoom-in process were always set to $L_1$ = 120 nm, $L_2$ = 80 nm. The L-value during the tracking measurement was set according to the signal-to-background ratio and the movement of the emitter. We set a waiting time of 5 µs between exposures to account for the time needed to switch the position of the intensity minimum with the phase modulator. The waiting time between measurements along subsequent axes was kept at 15 µs to account for the switching times of the acousto-optical deflectors.

**1.5. Diffusion: Theoretical background**

Diffusion at the single particle level is described by

$$D(t) = \frac{\mathrm{MSD}(t)}{2n_d t} \quad (1)$$

as introduced in the main text.

Here we describe how MSD(t) is obtained from a single particle trajectory and how the measurement itself affects it. We will also describe the two different types of diffusion observed in this study. Given a time trace with observations $\vec{r}_i$ interrupted by blinking events, one can calculate the mean squared displacement (MSD) for a given time delay $n\delta_t$

$$\mathrm{MSD}(n\delta_t) = \frac{1}{\sum_{i=1}^{N-n} a_i a_{i+n}} \sum_{i=1}^{N-n} (\vec{r}_{i+n} - \vec{r}_i)^2 a_i a_{i+n} \quad (2)$$

Here, $\delta_t$ is the time between two observations and $a_i = 1$ if the emitter is on at sample $i$ and $a_i = 0$ otherwise. $N$ is the total number of measurements and $n$ is the time difference between two measurements divided by $\delta_t$. For freely diffusing particles, the MSD increases linearly with time, i.e., the diffusion coefficient $D$ becomes independent of time. Therefore, Eq. 1 is simplified to

$$\mathrm{MSD}_{\mathrm{free}}(n\delta_t) = 2n_d D n\delta_t. \quad (3)$$

A measured MSD of a freely diffusing particle will deviate from this formula due to artifacts caused by the finite amount of time it takes to measure the continuously changing position of the particle. These artifacts are accounted for in the equation below (Ref. 2):

$$\mathrm{MSD}_{\mathrm{free}}(n\delta_t) = 2n_d D\delta_t(n - 2R) + 2n_d\Delta^2. \quad (4)$$

Here, $n_d$ is the number of dimensions in which the diffusion takes place, and $\Delta$ is the dynamic localization uncertainty resulting from the system's localization precision and spread of the fluorescence signal due to the emitter's movement. $R$ is a factor that accounts for the motion blur caused by the particle's movement during the time of measurement and is defined as (Ref. 3):

$$R = \frac{1}{\Delta_t}\int_0^{\Delta_t} S(t)[1 - S(t)]dt \quad (5)$$

where $S(t)$ is the integrated illumination up to time $t$ within a single measurement. For homogeneous illumination one obtains $R = 1/6$. The net effect of the two factors introduced in Eq.-4 is a vertical shift of the measured MSD values compared to the ground truth values. This behavior is also observed in Monte Carlo simulations, as shown in Fig. S3. If the stochastic movement of an object is hindered in any way, the diffusion properties of the particle may differ from the behavior observed in free diffusion. For example, if the particle is spatially confined, the diffusion is characterized by (Ref. 4)

$$\mathrm{MSD}_{\mathrm{conf}}(n\delta_t) = A[1 - \exp\left(-n\delta_t/\tau\right)] \quad (6)$$

where $A$ is the average distance between two randomly chosen points within the confinement region and $\tau$ is the equilibration time. For a 3D sphere with diameter $L$, $A \approx 0.26L^2$ and for a cube with edge length $L$, $A \approx 0.44L^2$. The equilibration time is given as a characteristic length scale of the confinement region divided by the diffusion coefficient inside the region. Like the parameter $A$, this characteristic length scale depends on the shape of the region. Since an analytical expression for $\tau$ could not be found, it was estimated from fits to Monte Carlo simulations of confined diffusion (according to SI Section 1.5.). Assuming spherical confinement regions, the inputs to the simulation could be reproduced in the fits when $\tau \approx 2A/D$. Plugging these parameters into Eq. 6 yields

$$\mathrm{MSD}_{\mathrm{conf}}(n\delta_t) = 0.26L^2\left[1 - \exp\left(-\frac{n\delta_t D}{0.52L^2}\right)\right] \quad (7)$$

Another type of diffusion occurs when the confinement regions are partially permeable, leading to a decoupling of diffusion on short and long timescales. This behavior is observed because the particle undergoes free diffusion on a short timescale, but encounters boundaries on a longer timescale that slow the diffusion. This type of diffusion is called hop diffusion and is characterized by a linear combination of free diffusion (Eq. 3) and confined diffusion (Eq. 7) (Refs. 2, 5)

$$\mathrm{MSD_{hop}}(n\delta_t) = \mathrm{MSD_{free}}(n\delta_t) + \frac{D_\mu - D_M}{D_\mu}\mathrm{MSD_{conf}}(n\delta_t)$$

$$= 2n_d n\delta_t \left\{D_M + \frac{D_\mu - D_M}{D_\mu}\frac{0.26L_{\mathrm{hop}}^2}{n\delta_t}\left[1 - \exp\left(-\frac{2n_d n\delta_t D_\mu}{0.52L^2}\right)\right]\right\} \quad (8)$$

Here, $D_\mu$ is the short-term diffusion coefficient and $D_M$ is the long-term diffusion coefficient. They are separated by a length scale $L_{\mathrm{hop}}$. Taking into account the measurement artifacts discussed above, Eq. 8 becomes

$$\mathrm{MSD_{hop}}(n\delta_t) = 2n_d \left\{D_M + \frac{D_\mu - D_M}{D_\mu}\frac{0.26L_{\mathrm{hop}}^2}{2n_d n\delta_t}\left[1 - \exp\left(-\frac{2n_d n\delta_t D_\mu}{0.52L^2}\right)\right]\right\}\delta_t(n - 2R) + 2n_d\Delta^2. \quad (9)$$

**1.6. Diffusion simulations and analysis routine**

To quantify the results of the diffusion measurements, an analysis routine was developed to determine the type and magnitude of the observed diffusion. This analysis routine was first thoroughly tested on Monte Carlo simulations to determine the limits of its validity.

*1.6.1. Simulation*

The simulations were performed with a custom written MATLAB code. The diffusion was simulated as a Wiener process in a 3D space divided into compartments by Voronoi tessellation. The input parameters to the simulation are the diffusion coefficient $D$, the radius of a Voronoi cell $L_{\mathrm{hop}}$, and a probability of transition to a neighboring cell $p_{\mathrm{hop}}$. Furthermore, the number of sampling points to be simulated and the time between these points are given to the simulation. In a first step, Voronoi cells are randomly distributed in space. Then, the center of a random Voronoi cell is taken as the starting point for the diffusion process. A single time step is taken by randomly sampling from a normal distribution with mean 0 and standard deviation according to the mean distance traveled in each spatial direction given by $D$ and the time between sampling points. A step is discarded with probability $1 - p_{\mathrm{hop}}$ if the final position is in a different cell than the position at the start of the step. Thus, for $p_{\mathrm{hop}} = 1$, one obtains free diffusion, and for $p_{\mathrm{hop}} < 1$, the Wiener process becomes increasingly obstructed, leading to hop diffusion. Next, the simulated position is linearly interpolated to increase the temporal precision so that a simulated MINFLUX measurement performed on this trace can take into account the movement of the measured object during each exposure while also providing the same temporal precision as the simulated trace. In the next step, a simulated 3D 4Pi MINFLUX measurement is performed on the interpolated trace. The input parameters of this simulation are the spatial width and offset of the illumination function, the exposure time per illumination and the $L$-value of the MINFLUX pattern. First, the input trace is transformed into the measurement coordinate system. This step is performed to provide the MINFLUX measurement routine with the 'ground truth' position of the measured object in the coordinate system in which the measurement is performed. The simulated measurement is performed along three axes with three exposures each and an L = 80 nm. For each single exposure, several detected photons are sampled from a Poisson distribution with its expectation value according to the simulated illumination function and the position of the measured object. For each cycle of nine exposures (3 per dimension), the position of the illumination pattern is adjusted according to the same localization algorithm applied by the FPGA during a real measurement. Finally, the entire trace is transformed back into the laboratory coordinate system by performing the inverse of the previous transformation. Examples of such simulated traces are shown in Fig. S3 for free diffusion ($p_{\mathrm{hop}} = 1$) and for hop diffusion ($p_{\mathrm{hop}} = 0.005$).

*1.6.2. Data analysis pipeline*

The goal of the data analysis pipeline is to calculate the MSD of a given trace. It is used to evaluate both measured and simulated traces. First, each trace is sliced into several sub-traces covering a time window of $n$ times the sampling time $\delta_t$. This step is taken to ensure uniform statistical sampling for each trace. A sub-trace is considered for further evaluation if at least 90% of its values meet the filter criteria. These are a minimum SBR of 1 per dimension and a positive curvature

$$c = 2\frac{n_+ + n_- - 2n_0}{L^2}. \quad (10)$$

The MSD is then calculated for each of the sub-traces according to Eq. 2.

*1.6.3. Data fitting*

Each MSD is subjected to a fitting routine to assess whether the measured object experienced free diffusion or hop diffusion and to determine the diffusion coefficient. The fitting process and differentiation between different diffusion modes follows a similar path as described in Ref. 2 with the main difference that the MSD curves are fitted instead of the diffusion coefficient curves. Fitting is performed only to the first data points of the MSD, as statistical errors dominate for larger time lags. In Ref. 5 it was suggested that the diffusion coefficient is best estimated by a linear fit to only the first few data points of the MSD. At the same time, other publications state that up to a quarter of the data points can be used (Ref. 6). For the analysis performed here, 20% of the data points have been used, as the aim is to distinguish between free and hop diffusion, which only becomes apparent at longer time lags. The fits are performed using the MATLAB function *fmincon*, which minimizes the sum of squared residuals (SSR) of the regression on the observed data. The fit functions are given in Eqs. 4 and 9 with $n_d = 3$ and $R$ depending on the type of data. For simulated data $\Delta = 0$ and $R = 0$, for simulated measured data $R = 1/6$ and for real measured data, $R \sim 0.15$ due to off-times during a single MINFLUX measurement caused by switching the illumination beam and the phase of the beam. For the fit to Eq. 9, the additional constraint $5D_M - D_\mu < 0$ is introduced to better distinguish free diffusion from hop diffusion. The values to which the fit is performed are sampled logarithmically to ensure that values for short time separations are weighted more heavily (Ref. 2) as they have higher statistical significance. Each fit is performed a predetermined number of times, using new randomized initial parameters for each run to avoid convergence to a local minimum. The two fits obtained are compared according to their Bayesian Information Criterion (BIC)

$$\mathrm{BIC} = n \ln\left(\frac{SSR}{n}\right) + k \ln(n).$$

Here, $n$ is the number of data points and $k$ is the number of fit parameters, i.e. $k = 2$ for free diffusion (Eq. 4) and $k = 4$ for hop diffusion (Eq. 9). The model with the smallest BIC is more likely and is used to classify the data as hop or free diffusion. Once the data are evaluated, all traces categorized as free diffusion are averaged and a fit is performed to obtain an ensemble estimate of the diffusion. The same is done for data categorized as hop diffusion.

*1.6.4. Test of analysis with simulated data*

To quantify the validity of the analysis routine, it was tested on a set of simulated traces. The simulated traces were performed with a photon emission rate and SBR similar to those observed in the experiments. The results of these tests are summarized in Tables S1 and S2. The data were evaluated for three different values of $n$, corresponding to lengths of 45 ms, 90 ms, and 180 ms, respectively. Table S1 shows the results for the simulation of traces undergoing free diffusion with a diffusion coefficient of $D = 10\,000\, nm^2/s$. The algorithm was able to correctly predict free diffusion for 85%-94% of the traces from the simulated ground truth for all tested values of $n$. The diffusion coefficient estimate is up to 20% larger than the ground truth value, and the misattributed result for hop diffusion gives an underestimate for macro diffusion and an overestimate for micro diffusion. 71%-84% of the traces from the simulated measurement could be correctly attributed to free diffusion. The diffusion coefficient was estimated to be within 20% of the ground truth for all values of $n$. Furthermore, the dynamic localization uncertainty is estimated to be about 6-7 nm, which is approximately the distance the measured object travels in a single time frame. The misclassified traces show too low macro diffusion and too high micro diffusion. In summary, the predicted values from the simulation and the simulated measurement are similar but different from the ground truth. Therefore, possible deviations from the expected behavior in the measurement are due to the analysis routine and not to the measurement process. The deviation of the diffusion coefficient for the simulations attributed to free diffusion is caused by traces with lower diffusion coefficients being incorrectly attributed to hop diffusion. The reason for the increasing misclassification for larger $n$ remains unknown. Table S2 shows the results for the simulation of traces undergoing hop diffusion with a diffusion constant of $D = 100\,000\, nm^2/s$, a compartment size of $L_{\text{hop}} = 30\, nm$ and a hopping probability of $p_{\text{hop}} = 0.001$. Hop diffusion could be correctly attributed to 29%-79% of the simulated traces, depending on the evaluated time period $n\delta_t$. The estimated value for $D_\mu$ is up to 20% larger than expected from the simulation input parameters and varies by about 10% depending on $n$. The estimated value for $D_M$ varies by almost a factor of two. Traces from simulated measurements could be correctly attributed for over 32%-88% of the traces. The values for $D_M$ and $D_\mu$ are similar to the simulated ground truth results. The value for $L_{\text{hop}}$ is up to twice as large as the ground truth value. Again, the analysis of simulated measurements and simulations of the ground truth show similar behavior. Therefore, the deviations are not due to the measurement process but to the analysis routine. The reason for the increase of $D_M$ and $D_\mu$ for smaller $n$ is caused by the increasing misclassification for the respective $n$. The reason for the misclassification remains unknown, but the result obtained with this simulation shows that the diffusion coefficients obtained from the analysis of the measured data are only accurate up to a factor of two. Overall, the analysis outlined above demonstrates the validity of the data fitting employed for traces undergoing hop diffusion in the sense that most traces are correctly categorized. In addition, the correct value for the diffusion coefficient in the case of free diffusion is obtained up to a maximum deviation of 20% (giving a quantitative result), and a separation of diffusion scales can be seen for hop diffusion, since $D_\mu/D_M \gg 5$ (giving a qualitative result).

#### 2. Precision limit in MINFLUX nanoscopy

A parabola can approximate the local minimum of the excitation profile. The fluorescence response of an emitter is then given by $I(x) = b + c\,(x - x_e)^2$, where b is the offset, c is the curvature of the parabola, and $x_e$ is the position of the emitter relative to the intensity minimum. Assuming that the intensity is below the saturation intensity of the illuminated emitter, it will emit a number of fluorescence photons proportional to the illumination intensity $N \propto I(x_e)$. Therefore, information about the position of the emitter $x_e$ can be extracted from the number of photons collected during the measurement. A measurement of the emitted photons for a single exposure is not sufficient to determine the position of the emitter, since the parameters b and c are unknown because they depend on a variety of parameters, such as the laser intensity, the background in the sample, and the quantum yield of the fluorophore, to name a few. To extract all three parameters, each MINFLUX measurement consists of a series of three exposures at equidistant positions $x = (-\frac{L}{2}, 0, \frac{L}{2})$ yielding the photon counts

$n_- = b(x_e - \frac{L}{2})^2 + a$

$n_0 = bx_e{}^2 + a$

$n_+ = b(x_e + \frac{L}{2})^2 + a$

This system of equations can be solved for the emitter position and for the uncertainty in the position. For a centered illumination pattern, the uncertainty is given by

$$\sigma = \frac{L}{4\sqrt{N}} \qquad (11)$$

Where $N = n_+ + n_- + n_0$ is the total number of photons detected. For non-zero background the uncertainty becomes

$$\sigma = \frac{L}{4\sqrt{N}}\sqrt{1 + \frac{5}{2\mathrm{SBR}} + \frac{3}{2\mathrm{SBR}^2}} \qquad (12)$$

Where $\mathrm{SBR} := \frac{cL^2}{4b}$ being the signal-to-background ratio. Eq. 12 shows that the localization precision cannot be improved indefinitely by choosing a small L in a measurement with a non-zero background, because the SBR decreases quadratically with smaller L. It has been shown (Ref. 7) that a minimally achievable localization precision of

$$\sigma_{\min} = \sqrt{\frac{b}{cN}} \qquad (13)$$

can be achieved with a minimal L where SBR = $\sqrt{3/2}$. Therefore, a MINFLUX illumination scheme must be optimized to have a small intensity minimum b and a high curvature c.

#### 3. Precision limit in 3D-4Pi MINFLUX

The local intensity minimum in 4Pi MINFLUX is generated by the destructive interference of two counterpropagating laser beams entering the sample through two opposing objectives. This is in contrast to the case of a single objective, where all beams are directed at the sample from a single side. The formation of the focal intensity distribution can be understood as the interference of plane waves, resulting in an intensity distribution of

$$I(x) \propto \sin^2\left(\frac{2\pi n \sin(\varphi) x}{\lambda_0}\right) \xrightarrow{Taylor} \frac{4\pi^2 n^2 \sin^2(\varphi)}{{\lambda_0}^2} x^2 := \frac{\pi^2}{\Lambda^2} x^2 \equiv c x^2 \tag{14}$$

Here, $\varphi$ is half the angle between the two interfering waves and $\lambda = \lambda_0/n$ is the wavelength in the medium. Furthermore,

$$\Lambda := \frac{\lambda_0}{2n\sin(\varphi)} \tag{15}$$

is the wavelength of the interferometrically generated illumination profile. The Taylor expansion around the intensity minimum shows that the curvature $c \propto 1/\Lambda^2$. Substituting this expression into Eq. 13 shows that $\sigma_{min} \propto \Lambda$. Therefore, the achievable localization precision is proportional to the wavelength of the illumination profile.

Due to the counterpropagating waves in 4Pi, $\varphi$ = 90°, resulting in the smallest possible illumination profile wavelength for a fixed excitation wavelength. To perform 3D measurements in a 4Pi configuration, the counterpropagating beams must be tilted with respect to the optical axis so that measurements can be performed along oblique axes while still utilizing the same optimal curvature as for measurements along the optical axis.

If the axes form a pattern where they are equidistantly spaced on a circle of radius r around the optical axis (Fig. 1B), an analytical expression for the coordinate transformation and the resulting localization precision can be obtained. A detailed derivation can be found in Ref. 8. Only the most important results are presented here.

An oblique axis can be addressed by shifting the beams from the optical axis so that the shift of the beam through one objective is the point reflection of the shift of the beam through the second objective, i.e. $(\Delta x, \Delta y)$ becomes $(-\Delta x, -\Delta y)$. For a lateral displacement of r from the optical axis in the back focal plane, the new axis forms an angle of

$$\delta = \arcsin\left(\frac{r}{fn}\right) \tag{16}$$

with the optical axis. Where f is the focal length of each objective and n is the refractive index of the immersion medium. At an angle of $\delta = 54.7°$, the three beams form a Cartesian coordinate system, i.e. the angle between each pair of axes is equal to 90°. In practice, the angle between the beam axes is less than 90° due to the limited NA of the objective lenses. The shearing angle $\beta$ is a measure of how much the coordinate system spanned by the three oblique axes is sheared relative to a Cartesian coordinate system. The relationship between $\delta$ and $\beta$ is:

$$\beta = \arctan\left(\frac{4-3\sqrt{1-\cos(-4\delta)}}{12\cos(-\delta)^2-8}\right) \tag{17}$$

The angles $\delta$ and $\beta$ are given in Fig. S4A as a function of the radius of the beam pattern in the back focal plane. A decrease in the pattern radius results in an approximately linear decrease in the angle of the individual beams with the optical axis. Since most pattern radii result in non-Cartesian coordinate systems, a coordinate transformation to the Cartesian laboratory system must be performed to obtain the true 3D position of a measured object. According to the transformation described in Ref. 8, uncertainties in the beam system propagate to the laboratory system. Assuming that the uncertainty along all three measurement axes is equal to $\sigma_{\mathrm{beam}}$, the uncertainty in the laboratory system is given by:

$$\sigma_{\mathrm{x}} = \sigma_{\mathrm{y}} = \sigma_{\mathrm{beam}}\sqrt{-\frac{\cos(\beta)\sqrt{1+2\tan(\beta)}}{\sin(2\beta)-1}} := \sigma_{\mathrm{beam}} T_{\mathrm{lat}}(\beta) \tag{18}$$

$$\sigma_{\mathrm{z}} = \sigma_{\mathrm{beam}} \frac{\cos(\beta)\sqrt{1+2\tan(\beta)}}{(\cos\beta + 2\sin\beta)} := \sigma_{\mathrm{beam}} T_{\mathrm{ax}}(\beta) \tag{19}$$

Here, $T_{\mathrm{lat}}$ and $T_{\mathrm{ax}}$ are the factors influencing the localization precision in the laboratory system based on the geometry of the beam system. These relationships are shown in Fig. S4B. For a shearing angle of $\beta = 0°$, all axes have the same $\sigma$, which is equal to $\sigma_{\mathrm{beam}}$. The larger $\beta$, the more the beam system is sheared towards the optical axis resulting in a smaller $\sigma_z$ while $\sigma_{x/y}$ are increased. The 3D localization precision is expressed by the third root of the detection volume, which is given by

$$\sigma_{3D} = \sqrt[3]{\sigma_x \sigma_y \sigma_z} \tag{20}$$

In general, the detection volume is optimal when $\beta = 0$.

In addition to the geometry of the beam system, the size of the individual beams also influences the maximum achievable localization precision, since the finite full width at half maximum (FWHM) of the beams leads to an effectively larger illumination profile wavelength $\tilde{\Lambda} := \Lambda\alpha$. The magnitude of $\alpha$ for a given FWHM has been calculated numerically and is shown in Fig. S4C. This correction arises due to the coherent summation of differently oriented plane waves in the focus.

In total,

$$\sigma_{x/y} \propto \Lambda(\lambda_0, n) * \alpha(\mathrm{FWHM}) * T_{\mathrm{lat}}(r, f, n)$$
$$\sigma_z \propto \Lambda(\lambda_0, n) * \alpha(\mathrm{FWHM}) * T_{\mathrm{ax}}(r, f, n)$$

i.e., the achievable localization precision depends on sample-dependent parameters such as the excitation wavelength, the focal length of the objectives, and the refractive index of the sample. On the other hand, it depends on the FWHM of the beams and their placement in the back focal plane, which can be optimized to achieve the best localization precision in a 4Pi configuration. In summary, to achieve optimal precision, the radius of the beam pattern in the focal plane must be chosen as large as possible so that $T_{\mathrm{lat}}$ and $T_{\mathrm{ax}}$ are minimized. In addition, the individual beams must be chosen as small as possible to keep $\alpha$ small. Keeping these two requirements in mind, one can optimize a 4Pi configuration to obtain the optimal $\sigma_{\mathrm{min}}$ with respect to the curvature c. The background factor b is not considered here because simulations show that it is zero, regardless of the specific illumination configuration.

##### 3.1. Comparison between single objective and dual objective MINFLUX

The illumination geometries referred to in this section are shown in Fig. S5.

In a single-objective MINFLUX microscope, a 1D local intensity minimum along the x/y-axis can be obtained by interference of two beams impinging on the back focal plane of the objective. The two beams are displaced from the optical axis by a distance r along the x/y-axis and are polarized perpendicular to the axis of displacement. A local intensity minimum can be obtained at the center of the focal plane if the beams have a relative phase of $\pi$ to each other. This configuration has been described in Ref. 7 and is sketched in Fig. S5C. In contrast to the 4Pi implementation presented in this paper, the achievable single-photon efficiency $\varepsilon$ depends only on the physical wavelength of the illumination profile, as no coordinate transformation is required. Furthermore, the influence of the individual beam sizes is not as pronounced as in a 4Pi illumination scheme. Numerical simulations show that in a single-objective MINFLUX implementation $\alpha < 1.05$ for all FWHM. Therefore, we only consider its influence in a 4Pi configuration for the calculations in this section.

The expression for the wavelength of the lateral illumination profile in a single objective configuration can be obtained by observing that the angle $\delta$ from Eq. 16 is equal to half the angle between the two interfering waves. Substituting Eq. 16 into Eq. 15 gives the expression for the wavelength of the lateral illumination profile in a single objective configuration:

$$\Lambda = \frac{\lambda_0}{2} \frac{f}{r} \tag{21}$$

Here, $\lambda_0$ is the wavelength of the beams in vacuum, f is the focal length of the objective, and r is the distance of the individual beams from the optical axis in the back focal plane. It is also possible to generate an axial intensity minimum in a single objective MINFLUX microscope. This has already been done with a tophat phase mask (Ref. 9) and a spatial light modulator (Ref. 10). In analogy to the method presented above for lateral illumination intensity synthesis, an axial minimum generated by three interfering beams is presented and discussed. The beam configuration is shown in Fig. S5D. It is similar to the configuration for lateral MINFLUX measurements, but has a third beam propagating along the optical axis. This third beam has the same polarization as the two outer beams. A local intensity minimum is generated when the phase of central beam is shifted by $\pi$ with respect to the outer beams. There is no analytical expression for the wavelength of the axial illumination profile. However, it is on the order of one $\lambda_0$ to several $\lambda_0$ depending on the pattern radius and the beam's FWHM. For the comparison between 3D 4Pi and single objective MINFLUX shown in Fig. S6, it was assumed that oil immersion objectives with f = 2 mm and n = 1.52 are used. We performed the calculations for beams with an FWHM = 1.5 mm, similar to the beam size used in the experiment, and for a beam size of FWHM = 0.1 mm, corresponding to widefield illumination. For widefield illumination, a 4Pi microscope outperforms a single objective microscope in terms of lateral single-

photon efficiency regardless of the chosen pattern radius, although the influence of the coordinate transformation increases for small pattern radii.
For the illumination scheme with FWHM = 1.5 mm, the lateral single-photon efficiency obtained with a single objective microscope is better than in a 4Pi microscope for pattern radii between 1-2 mm (see Fig. S6A). This is due to the fact that $\alpha$ increases for larger beams in the 4Pi case. The axial single-photon efficiency is at least a factor of three better in a 4Pi microscope compared to a single objective microscope (see Fig. S6B). The 3D single-photon efficiency is calculated from the single-photon efficiency along each dimension according to Eq. 20 (see Fig. S6C).
For beams with an FWHM = 1.5 mm, the 3D single-photon efficiency of a 4Pi microscope is about 50-60% better than a single objective microscope for pattern radii around 2 mm. It should be noted that the calculations were performed with an n = 1.52, which is the refractive index of the sample medium. Therefore, the above comparison is not valid for samples with a refractive index close to water (n = 1.33), which is the case for most biological samples. For such a sample, one could still use the oil immersion objective in a single objective microscope. Therefore, the single-photon efficiency would not change. At the same time, the wavelength of the illumination profile in the 4Pi sample would be extended by a factor of $\Delta = 1.52/1.33 \approx 1.14$, resulting in a 14% decrease in performance.
Nevertheless, the single-photon efficiency obtained in a 4Pi configuration will still outperform the single-photon efficiency obtained in a single objective microscope by at least 35%. Up to this point, we have only considered the curvature **c** for the comparison between single objective and 4Pi MINFLUX. The second factor affecting the single-photon efficiency is the residual background intensity b. We have not discussed this because there is no fundamental reason for background sources such as detector dark counts or autofluorescence from within the sample to differ between a single objective and a 4Pi microscope. Furthermore, the residual intensity in the minimum is theoretically zero for all illumination schemes presented in this section. Nevertheless, the quality of the intensity minimum obtained in an experiment will never match the simulations due to residual misalignments and aberrations. The main factors influencing the intensity minimum are the polarization, intensity, position, angle, collimation, and coherence of the individual beams. These beam parameters have complex relationships with different parts of the microscope. In fact, a 4Pi microscope consists of many more components than a single objective microscope, including but not limited to a second objective. This makes efficient alignment more difficult and introduces more sources of aberration. Therefore, a good intensity minimum is easier to achieve in a single objective microscope than in a 4Pi microscope.

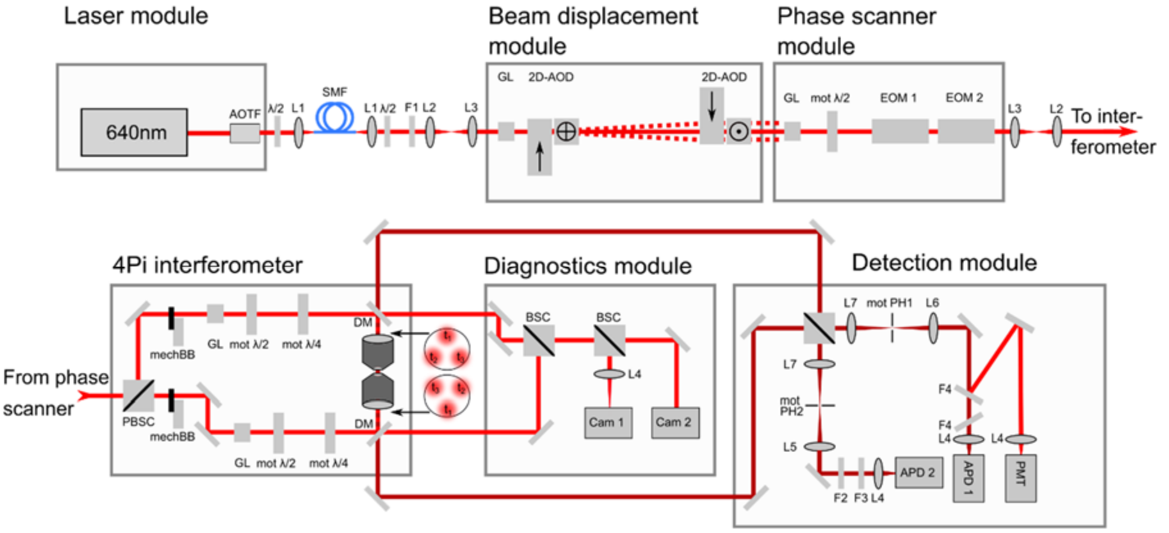


**Fig. S1. Sketch of the 3D-4Pi MINFLUX microscope**: The entire optical layout of the 3D-4Pi microscope, including all components described in SI Section 1.5. The beam displacement module is used to address the measurement axis, the phase scanner changes the local intensity minimum position along the beam propagation axis, and the beam is split and interfered in the 4Pi interferometer.

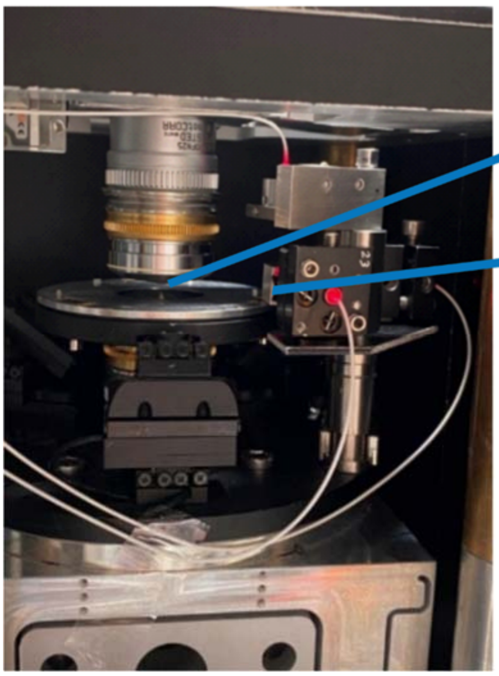

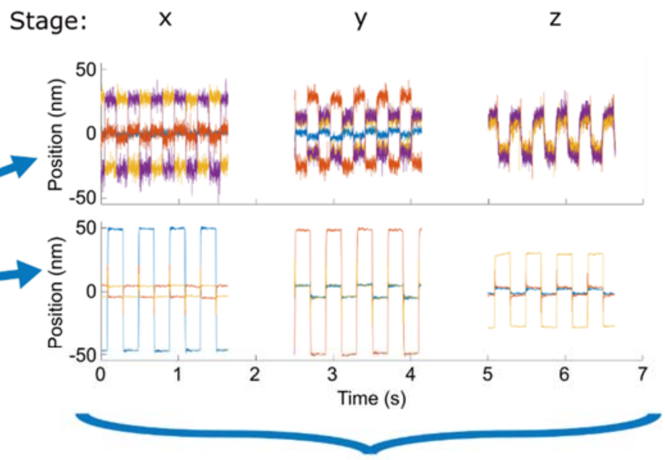


| | Beam 0 | Beam 1 | Beam 2 | Beam 3 |
|---|---|---|---|---|
| X | 0.12 | 0.15 | 1.02 | -0.90 |
| Y | -0.02 | 1.40 | -0.65 | -0.59 |
| Z | 0.59 | 0.61 | 0.57 | 0.30 |

**Fig. S2. Determining the coordinate transformation**: The picture shows the sample holder between the two objectives of the microscope. A mirrored cube is attached to the right side of the sample holder. The three white optical fibers come from the PicoScale device. They are attached to the custom-made construct on the right-hand side such that they can measure the position of the sample holder relative to the laboratory system. The positions from a MINFLUX measurement and a measurement of the stage position via the PicoScale during stage movements along the x, y, and z axis are shown in the plots on the right. The coordinate transformation from the measurement shown in the plots is given in the table at the bottom right. Beam 0 is a beam passing through the optical axis, while beams 1-3 are positioned as shown in Figure 1. A coordinate transformation from the measurements of beam 1-3 into the laboratory coordinate system is obtained from the same calibration measurement.

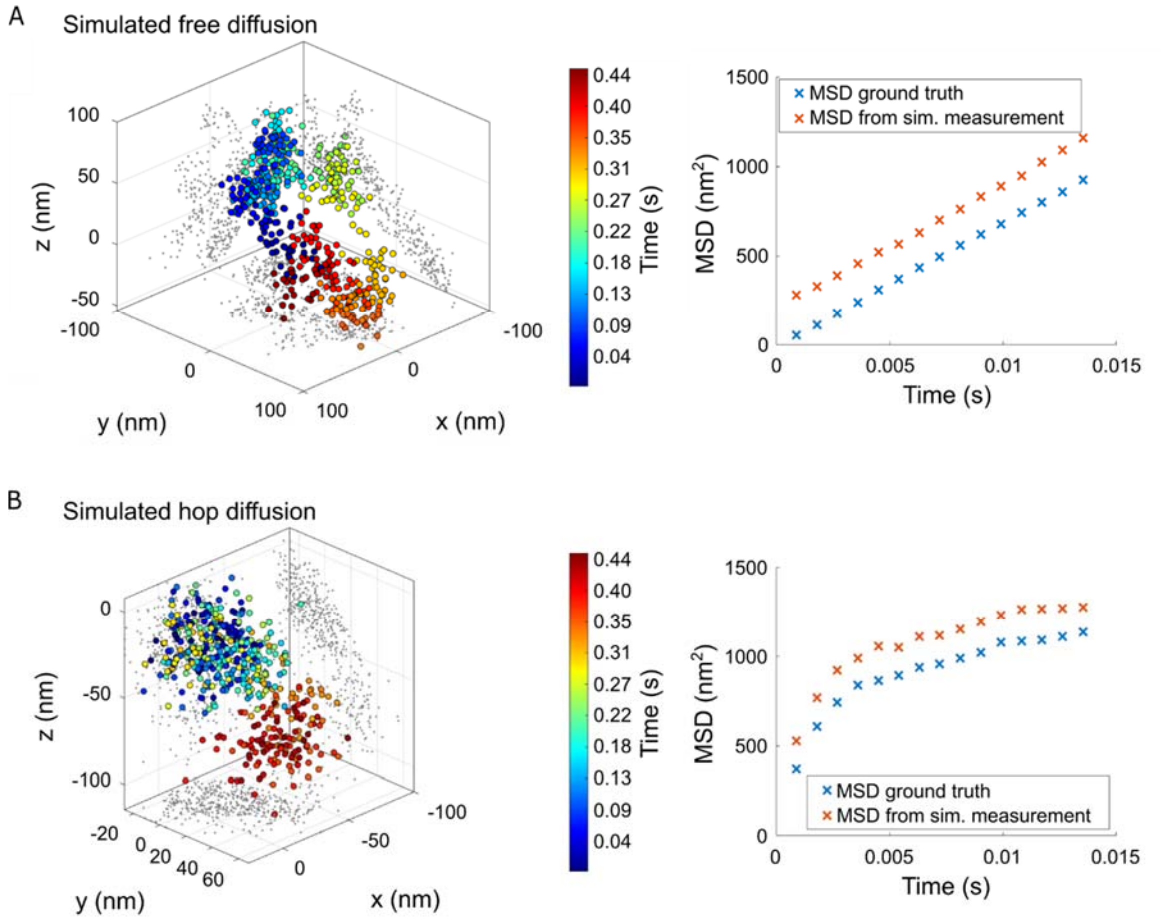


**Fig. S3. Simulated free and hop diffusion and corresponding MSD**: **A.** Left: Simulated measured positions of a particle undergoing free diffusion with a diffusion coefficient $\mathbf{D} = \mathbf{10\,000\,nm^2/s}$. Right: MSD calculated from the simulated ground truth positions and from the simulated measured positions. The simulated measurement shows an offset with respect to the ground truth due to the influence of the dynamic localization uncertainty (s. Eqs. 3 and 4). **B**. Left: Simulated measured positions of a particle undergoing hop diffusion with a diffusion coefficient $\mathbf{D} = \mathbf{100\,000\,nm^2/s}$, a compartment size $\mathbf{L_{hop}} = \mathbf{30\,nm}$ and a hopping probability $\mathbf{p_{hop}} = \mathbf{0.005}$. Right: MSD calculated from the simulated ground truth positions and the simulated measured positions. Again, the simulated measurement shows an offset with respect to the ground truth due to the influence of the dynamic localization uncertainty (s. Eqs. 8 and 9).

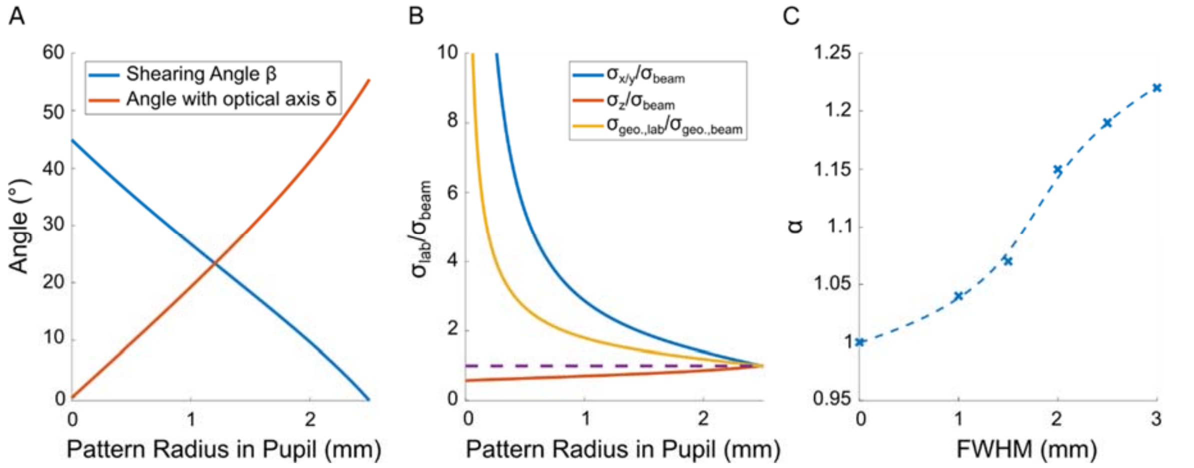


**Fig. S4 Factors influencing the *single-photon efficiency* in a 3D MINFLUX measurement with the 4Pi system**: **A.** Shearing angle β of the beam coordinate system relative to a Cartesian system and angle of the individual beams with the optical axis δ as given in Eqs. 17 and 16. The angles change approximately linearly with the pattern radius in the back focal plane. **B.** Ratio of σ in the laboratory system ($\boldsymbol{\sigma}_{\text{lab}}$) over σ in the beam system ($\boldsymbol{\sigma}_{\text{beam}}$) and ratio of the 3D detection volume ($\boldsymbol{\sigma}_{\text{geo}}$) in these two systems as a function of the pattern radius in the back focal plane. The dashed line is a reference with a ratio of one. For smaller pattern radii, the angle of the individual beams with the optical axis decreases, resulting in improved localization precision along the optical axis and decreased localization precision in the lateral plane. The detection volume becomes smallest when the beams form a Cartesian coordinate system. **C.** Numerical simulation of the increase in the illumination profile's wavelength($\widetilde{\boldsymbol{\Lambda}} := \boldsymbol{\Lambda\alpha}$) as a function of the beams' FWHM. Smaller beams lead to a smaller α, which leads to a higher curvature c. The dashed blue line is an interpolation to the values obtained from a numerical simulation of diffracting electromagnetic fields, shown as blue crosses.

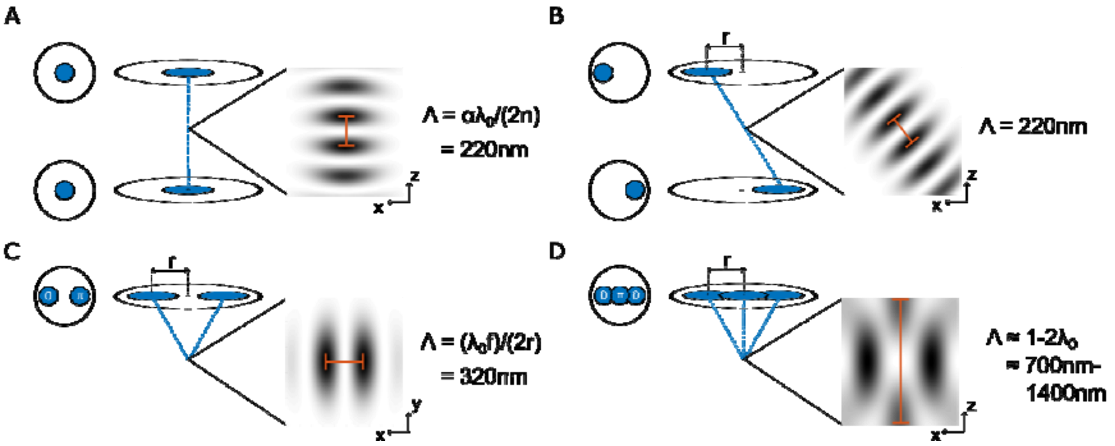


**Fig. S5. Illumination profiles for 1D-3D localizations in a single-objective and 4Pi microscope**: The values for the illumination profile's wavelengths $\Lambda$ were calculated for $\boldsymbol{\lambda_0 = 640\ \mathrm{nm}}$, $\mathbf{f = 2\ mm}$, $\mathbf{n = 1.52}$, $\mathbf{r = 2\ mm}$, and FWHM $= \boldsymbol{1.5\ mm}$. These values are typical for the measurements performed with the 4Pi **microscope** used in this work and for a single-objective MINFLUX microscope described in Ref. 7. The focal length is typical for Leica 100x oil immersion objectives. **A.** 1D 4Pi: Illumination configuration for axial measurements in a 4Pi microscope. The wavelength of the illumination profile is 220 nm. **B.** 3D-4Pi: One of the three beam pairs used to perform 3D measurements in a 4Pi microscope. As the beams are still counterpropagating, the wavelength of the illumination profile is the same as in the 1D case. **C.** 2D single objective: One of the two beam pairs used to perform 2D measurements in a single objective microscope. A phase difference of $\boldsymbol{\pi}$ between the two beams results in a central intensity minimum with a profile wavelength of 320 nm. **D.** 3D single-objective: A third illumination beam in the center of the two outer beams can be used to generate a local intensity minimum along the axial direction in a single objective MINFLUX microscope. The wavelength of the illumination profile is on the order of 1-2 $\boldsymbol{\lambda_0}$, depending on the pattern radius. For a radius of 2 mm, it is 720 nm.

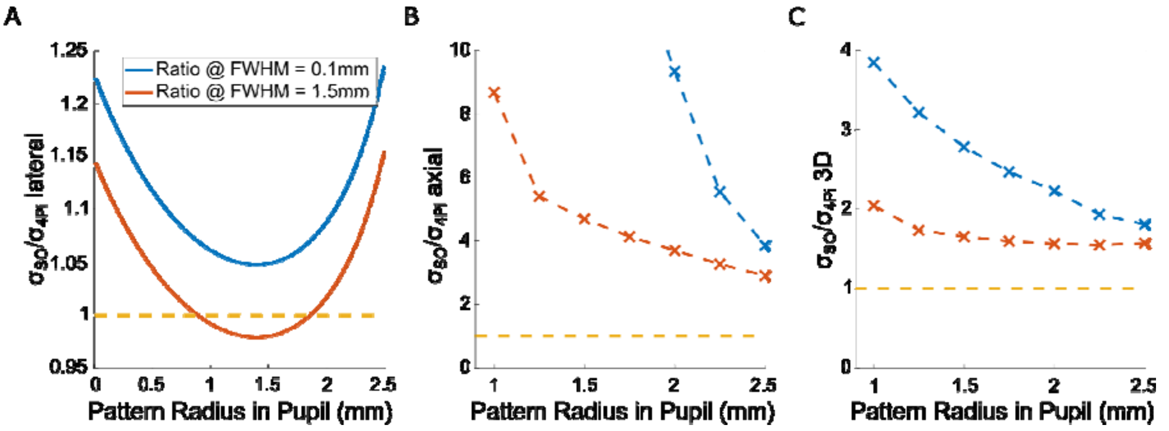


**Fig. S6. Single-photon efficiency in single objective vs. 4Pi MINFLUX**: All ratios shown were calculated for oil immersion objectives measuring a sample with an index of refraction of $\boldsymbol{n} = \mathbf{1.52}$. The blue curves show the values for beams with an FWHM $= \mathbf{0.1\ mm}$, and the red curves show the values for beams with an FWHM $= \mathbf{1.5\ mm}$. The yellow dashed line is introduced at a ratio of one as a guide for the eye. **A.** Ratio of single-photon efficiency for lateral localizations in a single objective microscope and a 4Pi microscope as a function of the pattern radius calculated according to Eqs. 15 and 21 with corresponding $\boldsymbol{\alpha}$ values. For beams with FWHM = 1.5 mm, the two microscopes have similar single-photon efficiencies, whereas for smaller beams, single-objective MINFLUX is outperformed by 4Pi MINFLUX. **B.** Numerically calculated ratio of single-photon efficiency for axial localizations in a single objective microscope and a 4Pi microscope as a function of pattern radius. A 4Pi configuration outperforms a single objective configuration by at least a factor of three. **C.** Ratio of single-photon efficiency for 3D measurements obtained from the calculated and simulated values shown in A and B, showing an increase in single-photon efficiency of about 50% for beams with FWHM $= \mathbf{1.5\ mm}$ and pattern radii around 2 mm.

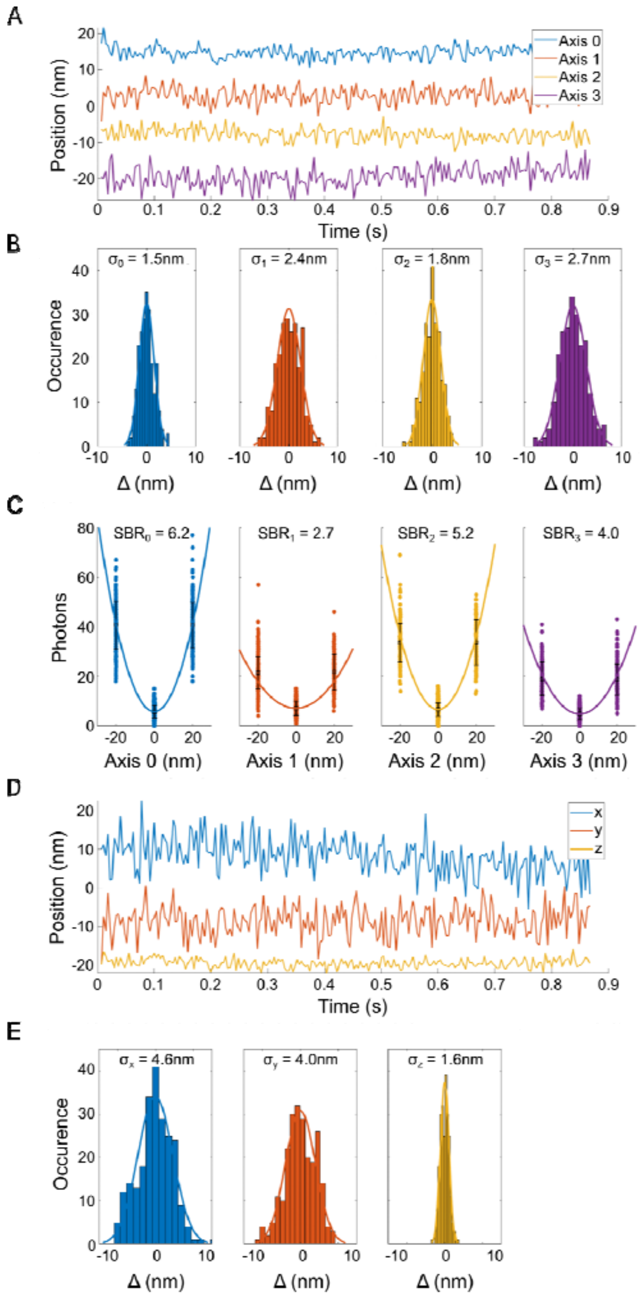


**Fig. S7. ATTO647N trace and statistics A.** ATTO647N trace measured along the optical axis (Axis 0) and the three outer axes spanning the 3D coordinate system of the MINFLUX measurement. **B.** Histograms of position differences between consecutive measurements obtained from the traces shown in A. **C.** Corresponding photon distributions and SBR along the four measured axes. **D.** Position of the emitter in the laboratory coordinate system obtained from a coordinate transformation of Axes 1-3. **E.** Histograms of position differences between consecutive measurements obtained from the traces shown in D, giving the localization precision in the laboratory coordinate system.

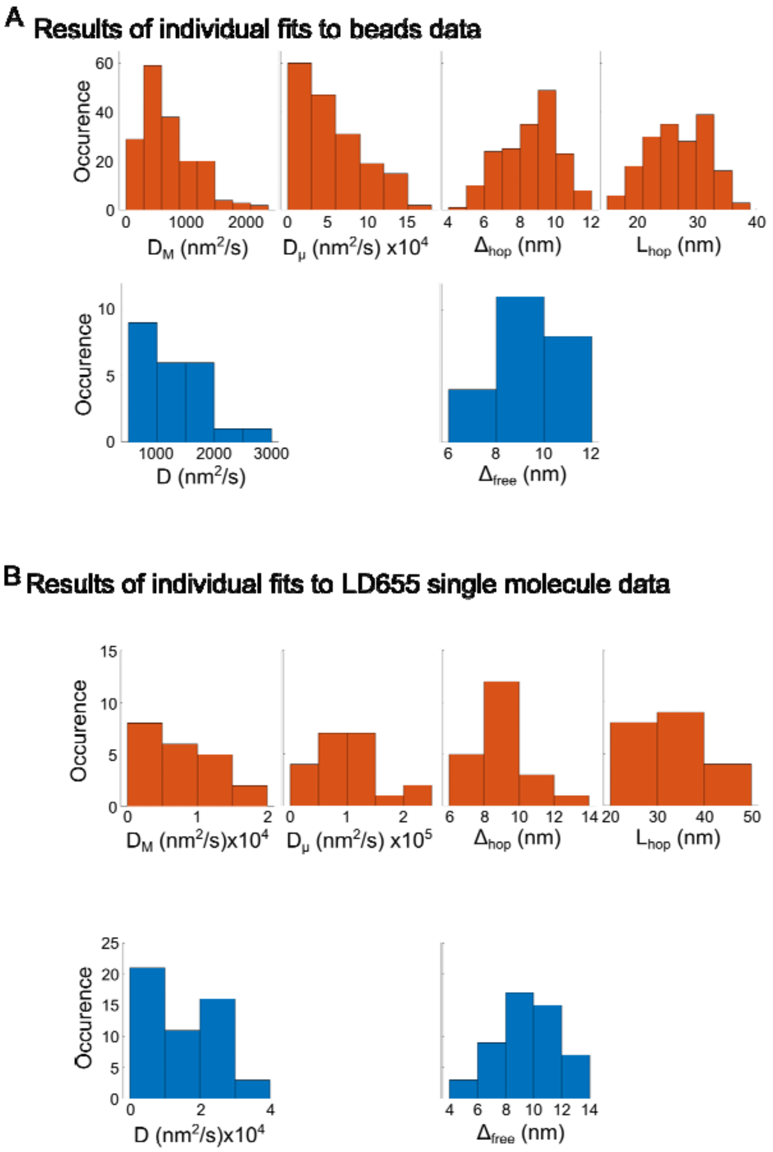


**Fig. S8. Results of individual fits to diffusion traces: A.** Histograms of fitted parameters for all single bead traces categorized as hop diffusion are shown in orange. Results for traces categorized as free diffusion are shown in blue. The histograms show the variance of the observed values due to different nanoenvironments. **B.** Histograms of fitted parameters for all single LD655 singe molecule traces categorized as hop diffusion are shown in orange. Results for traces categorized as free diffusion are shown in blue. The histograms show the variance of the observed values due to different nano-environments.

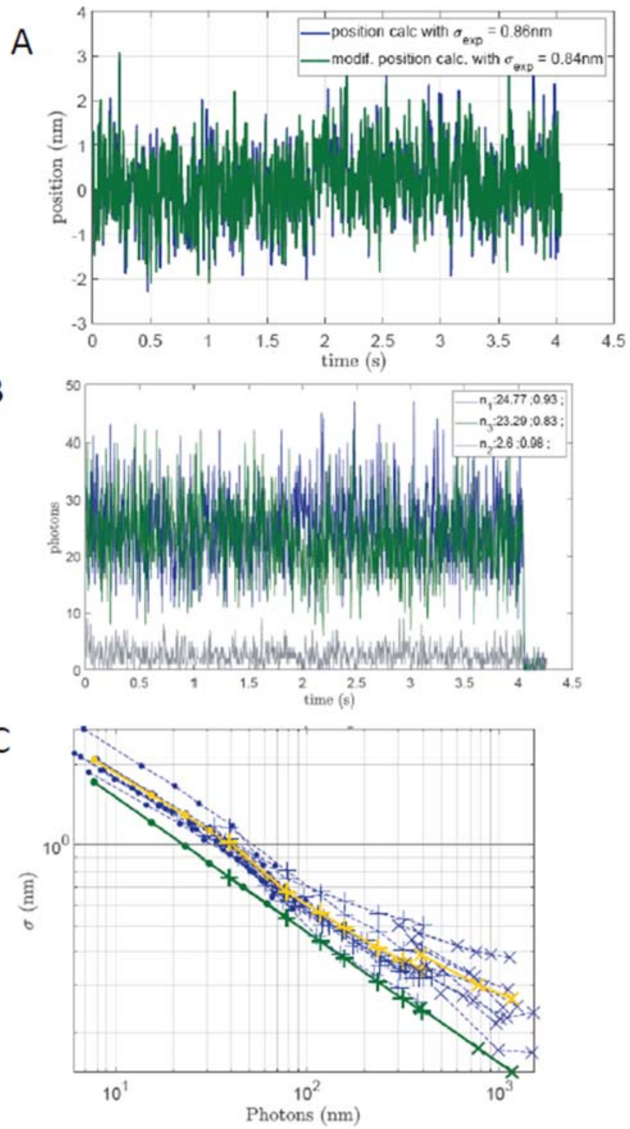


**Fig. S9. ATTO647N trace and statistics measured along the optical axis in an earlier stage of the experimental setup A.** Position trace of the emitter along the axial direction. **B.** Measured photon counts of the trace in **A**. **C.** Axial localization uncertainty for the statistics of emitters.

**Table S1. Fitting results for the simulation of free diffusion**: Free diffusion with a diffusion constant of $\mathrm{D} = 10\,000\ \mathrm{nm}^2/\mathrm{s}$ was simulated repeatedly at a sampling rate of 1.1 kHz. The resulting traces were evaluated for three different values of n (corresponding to 45 ms, 90 ms, and 180 ms) according to the analysis pipeline outlined in SI Section 1.5. The results of this analysis are shown here. The results that specify the diffusion mode and quantify the speed of diffusion are highlighted in blue.

| | | n | 50 | 100 | 200 |
|---|---|---|---|---|---|
| Simulation | Classification | free | 94% | 88% | 85% |
| | | hop | 6% | 12% | 15% |
| | Results free fit | $\boldsymbol{D}(\boldsymbol{nm}^2/\boldsymbol{s})$ | 10000 | 11000 | 11000 |
| | | $\Delta_{\text{free}}(\boldsymbol{nm})$ | 0 | 0 | 0 |
| | Results hop fit | $\boldsymbol{D}_{\boldsymbol{\mu}}(\boldsymbol{nm}^2/\boldsymbol{s})$ | 22000 | 25000 | 24000 |
| | | $\Delta_{\text{hop}}(\boldsymbol{nm})$ | 0 | 0 | 0 |
| | | $\boldsymbol{L}_{\text{hop}}(\boldsymbol{nm})$ | 17 | 26 | 37 |
| | | $\boldsymbol{D}_{\boldsymbol{M}}(\boldsymbol{nm}^2/\boldsymbol{s})$ | 4400 | 5000 | 4900 |
| Simulated measurement | Classification | free | **84%** | **78%** | **71%** |
| | | hop | 16% | 22% | 29% |
| | Results free fit | $\boldsymbol{D}(\boldsymbol{nm}^2/\boldsymbol{s})$ | **10000** | **11000** | **12000** |
| | | $\Delta_{\text{free}}(\boldsymbol{nm})$ | 6.6 | 6.7 | 6.5 |
| | Results hop fit | $\boldsymbol{D}_{\boldsymbol{\mu}}(\boldsymbol{nm}^2/\boldsymbol{s})$ | 44000 | 27000 | 29000 |
| | | $\Delta_{\text{hop}}(\boldsymbol{nm})$ | 6.0 | 6.4 | 6.1 |
| | | $\boldsymbol{L}_{\text{hop}}(\boldsymbol{nm})$ | 27 | 32 | 38 |
| | | $\boldsymbol{D}_{\boldsymbol{M}}(\boldsymbol{nm}^2/\boldsymbol{s})$ | 6600 | 5400 | 5900 |

**Table S2. Fitting results for the simulation of hop diffusion:** Confined diffusion with a diffusion constant of $\mathbf{D} = \mathbf{100\,000\ nm^2/s}$, a compartment size of $\mathbf{L_{hop}} = \mathbf{30\ nm}$ and a hopping probability of $\mathbf{p_{hop}} = \mathbf{0.001}$ was simulated repeatedly at a sampling rate of 1.1 kHz. The resulting traces were evaluated for three different values of n (corresponding to 45 ms, 90 ms, and 180 ms) according to the analysis pipeline outlined in SI Section 1.5. The results of this analysis are shown here. The results that specify the diffusion mode and quantify the speed of diffusion are highlighted in blue.

| | | n | 50 | 100 | 200 |
|---|---|---|---|---|---|
| Simulation | Classification | free | 71% | 38% | 21% |
| | | hop | 29% | 62% | 79% |
| | Results free fit | $D(nm^2/s)$ | 44000 | 52000 | 73000 |
| | | $\Delta_{free}(nm)$ | 7.1 | 7.0 | 1.9 |
| | Results hop fit | $D_\mu(nm^2/s)$ | 120000 | 120000 | 110000 |
| | | $\Delta_{hop}(nm)$ | 4.8 | 6.0 | 6.6 |
| | | $L_{hop}(nm)$ | 42 | 52 | 63 |
| | | $D_M(nm^2/s)$ | 24000 | 19000 | 14000 |
| Simulated measurement | Classification | free | 68% | 28% | 12% |
| | | hop | **32%** | **72%** | **88%** |
| | Results free fit | $D(nm^2/s)$ | 39000 | 47000 | 60000 |
| | | $\Delta_{free}(nm)$ | 8.8 | 8.8 | 7.0 |
| | Results hop fit | $D_\mu(nm^2/s)$ | **150000** | **120000** | **110000** |
| | | $\Delta_{hop}(nm)$ | 6.0 | 7.5 | 8.0 |
| | | $L_{hop}(nm)$ | 46 | 52 | 64 |
| | | $D_M(nm^2/s)$ | **27000** | **18000** | **14000** |
